# The genomics of trait combinations and their influence on adaptive divergence

**DOI:** 10.1101/2020.06.19.161539

**Authors:** McCall B. Calvert, Meredith M. Doellman, Jeffrey L. Feder, Glen R. Hood, Peter Meyers, Scott P. Egan, Thomas H.Q. Powell, Mary M. Glover, Cheyenne Tait, Hannes Schuler, Stewart H. Berlocher, James J. Smith, Patrik Nosil, Daniel A. Hahn, Gregory J. Ragland

## Abstract

Understanding rapid adaptation requires quantifying natural selection on traits and elucidating the genotype-phenotype relationship for those traits. However, recent studies have often failed to predict the direction of adaptive allelic variation in natural populations from laboratory studies. Here, we test for genomic signatures of genetic correlations to illustrate how multifarious, correlational selection may drive counterintuitive patterns of population divergence in the apple maggot fly, *Rhagoletis pomonella* (Diptera: Tephritidae). Apple-infesting populations with relatively early emerging adults have recently evolved from hawthorn-infesting populations consisting of relatively later emerging adults. Multiple studies have reported a paradoxical relationship between population differentiation and seasonal timing, as determined by the timing of diapause termination; alleles associated with late termination occur at higher frequencies in the earlier emerging apple-infesting populations compared to hawthorn-infesting populations. We present evidence that historical selection on diapause termination and another trait, initial diapause intensity, over geographic gradients generated genetic correlations between the traits in a direction antagonistic to contemporary selection on apple-infesting populations. Single nucleotide polymorphism in genomic regions of high linkage disequilibrium associated strongly with diapause termination and intensity, population divergence, geography, and evolutionary responses in laboratory selection experiments. These associations were consistent with geographically variable selection and with correlated evolutionary responses driving higher frequencies of late-associated alleles in the early emerging apple race. In contrast, loci associated only with diapause termination showed the expected pattern (more early-associated alleles in the apple race) in half of the population pairs. Our results suggest that selection on loci demonstrating antagonistic pleiotropy may often shape genomic footprints of rapid adaptation.

## INTRODUCTION

A common goal among evolutionary biologists is to evaluate the genetic basis of adaptation on ecological time scales (Futuyma, 2010; Stinchcombe & Hoekstra, 2007). Developing a comprehensive understanding of rapid adaptation requires knowledge of natural selection, the genotype-to-phenotype relationships for traits responding to selection, and standing genetic variation for those traits (Pfrender, 2012). The clearest descriptions of the genomic basis of adaptation have been made in systems where well-defined, directional selection consistently operates on traits controlled by a handful of loci of major effect (Etges, Oliveira, Noor, & Ritchie, 2010; F. C. Jones et al., 2012; M. R. Jones et al., 2018; Peichel & Marques, 2017; Soria-Carrasco et al., 2014; Via, Conte, Mason-Foley, & Mills, 2012). Selection experiments on specific traits have replicated natural genetic variation among diverging populations in several classic cases, including stickleback fish lateral plate development (*Ectodysplasin* gene; Barrett, Rogers, & Schluter, 2008; Jones et al., 2012), beak size in Darwin’s finches (*ALX1* haplotype; Lamichhaney et al., 2015), *Heliconius* butterfly wing color polymorphism (the *optix* genes; Reed et al., 2011), color polymorphisms in *Timema* stick insects (*Mel-Stripe* locus; Gompert et al., 2014; Nosil et al., 2018), and coat color in field mice (*agouti* gene; Barrett et al., 2019; Steiner, Weber, & Hoekstra, 2007). However, the association between trait variation and variation at focal loci typically includes substantial uncertainty and/or unexpected trends (Nosil et al., 2018; Rennison, Heilbron, Barrett, & Schluter, 2015). Depending on the context, this pattern likely results from incomplete knowledge on fluctuations in selection over time and space, the targets of natural selection, and imperfect understanding of genetic relationships among those targets and the rest of the genome.

Under polygenic models of adaptation, hundreds to thousands of genetic loci contribute to variation in a trait under selection and, therefore, the effect of any single locus is expected to be small (Boyle, Li, & Pritchard, 2017; Rockman, 2012). The weak signal of any given variant is difficult to detect, and response to selection can appear as genome-wide shifts in allele frequencies or decreases in nucleotide diversity suggestive of soft selective sweeps (Berg & Coop, 2014; Hermisson & Pennings, 2005). The ability to identify causal variants and architectures of adaptation is strengthened when genome wide association (GWA) studies for traits putatively under selection are combined with genome scans for divergent loci between populations (Brennan et al., 2018; Egan et al., 2015; Gompert et al., 2014; McGirr & Martin, 2017). This experimental design also allows for a deeper dissection of the potential genetic mechanisms mediating adaption, as it allows us to quantify the contribution of single ecological or evolutionary processes to adaptive variation in natural populations. Such approaches have often demonstrated a high degree of *genomic predictability*: a strong alignment between the loci underlying an adaptive, quantitative trait and loci contributing to population differentiation (Brennan et al., 2018; Fountain et al., 2016; Orsini, Spanier, & Meester, 2012; Soria-Carrasco et al., 2014; Troth, Puzey, Kim, Willis, & Kelly, 2018; Via et al., 2012). However, the sign of allele frequency change between divergent populations in nature does not always match expectations derived from laboratory observations. That is, some alleles that appear adaptive based on associations with a phenotype or a laboratory selection response occur at low frequencies in the natural population experiencing selection, as seen in Melissa blue butterflies (Chaturvedi et al., 2018), *Daphnia* water fleas (Orsini et al., 2012), pea aphids (Via et al., 2012), and Glanville fritillary butterflies (Fountain et al., 2016). One explanation is that different phase or linkage disequilibrium (LD) relationships may exist between the true causal variants and genotyped single nucleotide polymorphisms (SNPs) across different populations (Linnen, 2018). A natural extension of this hypothesis is that genomic predictability should increase with the proportion of the genome that is sequenced. However, whole genome sequence data from *Timema* stick insects did not reveal elevated levels of genomic predictability between different host-associated populations compared to other studies using reduced representation sequencing (Soria-Carrasco et al., 2014).

Additionally, a lack of genomic predictability (alignment between loci associated with an adaptative trait and loci associated with population differentiation) may be the result of incomplete knowledge of the multivariate trait combinations responding to natural selection. Fitness is generally determined by high-dimensional, multivariate (or multifarious) selection (Blows & Hoffmann, 2005; Orr, 1998; Saltz, Hessel, & Kelly, 2017). Thus, adaptation to novel environments can involve trait shifts along multiple ecological niche axes, as is the case with Lake Victoria cichlid species which vary in diet, parasite load, and visual system (male color and female preference) with water depth (Seehausen, 2009; Seehausen et al., 2008). Moreover, adaptation may be facilitated (or constrained) if adaptive traits are genetically correlated through mechanisms such as pleiotropy, high physical linkage, or structural variation (Via and West 2008; Feder et al. 2012). Several theoretical models predict that reduced recombination among combinations of alleles within co-adapted gene complexes should promote adaptation and speciation (Feder & Nosil, 2009; Kirkpatrick & Barton, 2006; Navarro & Barton, 2003), though few studies provide rigorous empirical tests (but, see Ayala et al., 2019; Coughlan & Willis, 2019; Ruegg, Anderson, Boone, Pouls, & Smith, 2014). Such tests are necessary to fully resolve the degree to which the genetic basis of trait correlations influences the predictability of evolution, and at least one study presents clear evidence for antagonistic pleiotropy among fitness related traits mediating the predictability of evolutionary response (Troth et al., 2018).

The apple maggot fly *Rhagoletis pomonella* (Diptera: Tephritidae) is a well-studied system with evidence for multifarious selection driving population divergence in phenotypes, yet observed, counterintuitive trends in the frequencies of alleles associated with those phenotypes (Feder, Hunt, & Bush, 1993; Michel et al., 2010; Ragland et al., 2017). The *R. pomonella* complex is a group of closely related, North American fruit-feeding species, each uniquely adapted to a different fruit host (Feder, Chilcote, & Bush, 1988; Xie et al., 2008). Fly larvae develop within this fruit during the summer or fall, exit the fruit to pupate in the soil, and then overwinter in diapause, a state of developmental arrest (Dean & Chapman, 1973). When diapause terminates the following summer, adults emerge, mate, and lay eggs in the host fruit (Dean & Chapman, 1973). Host associated-populations of *R. pomonella*, also termed host races, can evolve rapidly. Apple-infesting host races formed approximately 160 years ago (Bush, 1969; Walsh, 1867), are genetically differentiated from ancestral hawthorn-infesting populations (Feder et al., 1988; McPheron, Smith, & Berlocher, 1988), and have different host odor preferences (Linn et al., 2004, 2003). Hereafter, we refer to these as the apple and hawthorn races. Seasonal emergence time has also rapidly evolved; apple race flies emerge as adults earlier than hawthorn race flies at sympatric sites, synchronizing with the earlier fruiting time of apple trees (Doellman et al., 2019; Feder et al., 1994). The timing of adult emergence from overwintering pupae is developmentally controlled by the timing of pupal diapause termination (Fig 1a; Ragland et al. 2009, 2011). Past studies using allozyme markers, microsatellites, and genome-wide SNP markers all found loci strongly associated with diapause termination timing and with host race divergence (Feder et al., 1993; Michel et al., 2010; Ragland et al., 2017). However, these studies also found that the direction of allelic variation in the recently derived apple race was consistently opposite to the expected direction such that the earlier emerging apple race harbored higher frequencies of alleles associated with later diapause termination.

**Figure 1:**
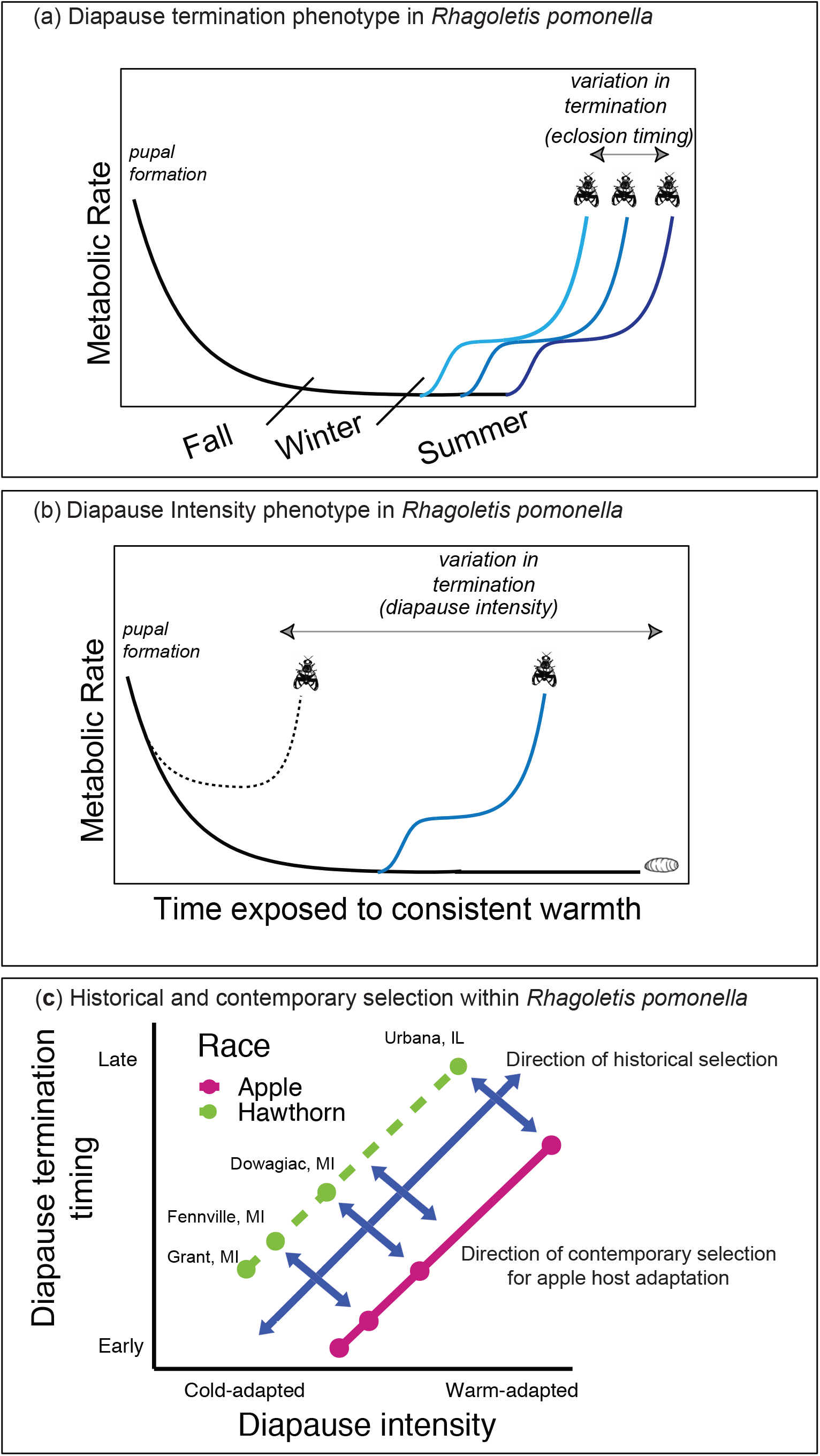
Conceptual diagram of historical and contemporary selection on *R. pomonella* diapause life history traits. (a) The timing of diapause termination, which controls adult emergence time, varies within and between host races (different colored trajectories) and is defined as the timing of the initial, rapid increase in metabolism that is followed by the characteristic, biphasic trajectory of post-diapause development (Ragland *et al.*, 2009). (b) Initial diapause intensity is measured as the propensity to forgo diapause and develop into an adult when exposed to artificially warm conditions simulating long pre-winter periods. This can occur if an individual does not enter diapause (dashed line, Non-Diapause [ND]) or if an individual prematurely terminates diapause (solid blue line, Shallow Diapause [SD]). (c) *R. pomonella* is a summer and fall breeding species because its life cycle tracks the availability of fruits that mature during the field season. In the Midwestern range of *R. pomonella*, growing seasons shorten and winters increase in intensity with increasing latitude. As a result, within the ancestral hawthorn race, phenotypic combinations of a more warm-adapted diapause intensity and late diapause termination for southern populations and more cold-adapted diapause intensity and early diapause termination for northern population are favored. Whereas, contemporary selection for the formation of the apple race favors a relatively warmer adapted diapause intensity and earlier diapause termination at a sympatric, geographic site. Here, contemporary selection is perpendicular to trait combinations geographically favored within the hawthorn-infesting host race.

As a summer and fall-breeding species, mean diapause termination time of geographic populations of *R. pomonella* increases with decreasing latitude to track later fruiting times of their fruiting host plants (Feder and Bush 1989; Dambroski and Feder 2007). This increase in diapause termination timing with decreasing latitude corresponds with an increase in a second diapause related trait (Dambroski & Feder, 2007), which we term diapause intensity. Diapause intensity is defined by an individual’s recalcitrance to pre-winter cues and may also evolve rapidly and contribute to host race divergence. Warm pre-winter conditions in the summer and fall can cause individuals to terminate diapause prior to winter (Dambroski & Feder, 2007; Prokopy, 1968), resulting in premature adult emergence and overwintering mortality (Fig. 1b). Individuals with a more intense diapause tend to remain in diapause despite warm temperatures (Dambroski & Feder, 2007; Egan et al., 2015). The cooccurring phenotypic clines for diapause termination and diapause intensity suggests that these traits may be correlated, leading us to hypothesize that selection for diapause intensity is ‘dragging along’ maladaptive variation in *R. pomonella* diapause termination timing through pleiotropy, LD, or a combination of the two (Fig. 1c). Thus, historic, correlational selection along geographic clines in the ancestral hawthorn race has favored combinations of late diapause termination timing and more intense diapause in the South and early diapause termination timing and less intense diapause in the North (Fig. 1c). In contrast, selection on the apple compared to the hawthorn race at sympatric sites favors relatively earlier diapause termination timing and more intense diapause in the apple race in response to earlier fruiting apple trees that also leads to warmer, longer prewinter periods (Egan et al., 2015; Feder, Roethele, Wlazlo, & Berlocher, 1997; Ragland et al., 2017). Therefore, host-associated selection favors trait combinations that have been historically disfavored by geographically variable selection in the ancestral host race, potentially affecting and constraining the recent evolution of host-associated traits.

We further hypothesized that historical, correlational selection in *R. pomonella* has favored increased LD between loci or an increase in pleiotropically acting loci promoting an association between earlier diapause termination timing and increased diapause intensity. Specifically, we test hypotheses about whether the two traits are genetically correlated, what processes may have generated that correlation, and how contemporary selection on correlated traits influences adaptation to novel hosts. To test these hypotheses, we combined new data on genotype-to-phenotype relationships for diapause intensity with published data on genotype-to-phenotype relationships for diapause termination, population genetic variation among host races and across geography, and genetic responses to laboratory selection experiments that mimic field seasonality (Table 1; Doellman et al., 2019, 2018; Egan et al., 2015; Ragland et al., 2017). We asked five questions about the influence of pleiotropy and/or LD on observed genetic differences between the *R. pomonella* host races: 1) Is there segregating genetic variation for diapause intensity in the ancestral, hawthorn race?; 2) Do genetic variants associated with diapause intensity also associate with host race divergence?; 3) Do genetic variants associated with diapause intensity also associate with diapause termination?; 4) Do genetic variants associated with diapause intensity also respond to laboratory selection on seasonality?; 5) Do genetic variants associated with diapause intensity and diapause termination vary geographically according to expectations based on historical, correlational selection?

**Table 1:** Description of the genomic datasets using RAD-seq generated in previous studies of *R. pomonella*.

| Experiment | Description | Reference |
| --- | --- | --- |
| Diapause termination GWA study | Extreme phenotyping analysis of genetic associations with diapause termination time in both host races. | Ragland et al., 2017 |
| Pre-winter selection | Both host races were exposed to a short (7-day) vs. a long (32-day) pre-winter period to simulate prewinter conditions typically experienced by the hawthorn and apple races in nature, respectively, followed by genotypic comparisons pre- and post-selection. The hawthorn race study included a genetic host race comparison at Grant, MI. | Egan et al., 2015 (hawthorn race)<br>Doellman et al., 2018 (apple race) |
| Geographic clines | Random samples from the apple and hawthorn races reared from field collected fruit at Fennville, MI, Dowagiac, MI, and Urbana, IL were compared against each other (including previous Grant, MI data) and within each site to assess geographic variation relative to host divergence. | Doellman et al., 2019 |

## 2. METHODS & MATERIALS

### 2.1 Fly collection and rearing

Standardized collection and rearing methods were used for the current study and to generate all other, previously published data included in the analyses (Egan et al., 2015). Adult flies were reared from field-collected fruit. Specifically, field-collected, infested fruit were held over wire mesh racks on top of plastic collecting trays which were checked for pupae a minimum of three times per week. For diapause termination timing and diapause intensity phenotyping, we cleared trays in the early evening and only used flies in the experiment that exited from fruit and formed puparia in the intervening period from early evening to the following morning to synchronize development. Collected pupae were placed in petri dishes with moist vermiculite and held for 10 days at 14:10 L:D and 21°C, then transferred to the desired experimental conditions. Unless otherwise noted, overwintering treatments for all experiments were conducted at 4°C and constant darkness.

### 2.2 Double digest restriction-site associated DNA sequencing (RAD-seq)

Genotypic data (SNPs) for the current study and for all other, previously published studies included in the analyses were generated using a common RAD-seq approach (Parchman et al., 2012). Generation of individually barcoded double digest restriction amplified DNA (ddRAD) libraries, *de novo* genome assembly of contigs, variant SNP calling, and allele frequency estimation were performed following Egan et al. (2015). Sequencing was performed on Illumina HiSeq platforms. Custom scripts and the Genome Analysis Toolkit (GATK version 2.5-2; McKenna et al. 2010) were used to estimate the genotype probabilities used as input for the analyses described below, which account for genotype uncertainty.

### 2.3 Previously published RAD-seq studies

#### 2.3.1 Geographic survey

The RAD-seq data comparing sympatric host races at multiple geographic sites from Doellman et al., (2019) were used to investigate how loci associated with diapause traits vary across geographic gradients in seasonality. In Doellman et al. (2019), adult flies from both apple and hawthorn races were reared from field collected fruit gathered at sympatric sites in Grant (lat., long. = 43.35 N, −85.9 W), Fennville (42.6 N, −86.15 W), and Dowagiac, Michigan, USA (41.88 N, −86.23 W), and Urbana, Illinois, USA (40.08 N, −88.19 W). These sites roughly follow a north–south cline in the Midwestern United States. Larvae that emerged from fruit and pupated were overwintered and reared to adulthood following the methods of Egan et al. (2015).

#### 2.3.2 Diapause termination GWA study

The RAD-seq data used to associate SNP genotypes with diapause termination phenotypes are from Ragland et al. (2017). Briefly, pupae from both host races sampled from Fennville, Michigan were reared from field-collected fruit as described above, overwintered for 16 weeks, then held at 21°C, 14:10 L:D until adult emergence from the pupae. The earliest and latest 4% of emerging flies in each host race were sequenced in an extreme phenotyping design (similar to bulked segregant analysis), wherein the strength of genetic association with diapause termination timing was determined by the magnitude of allele frequency differences between the bulks (i.e., groups of individuals from opposite tails of the emergence distribution; Michelmore, Paran, & Kesseli, 1991; Pool, 2016). Allele frequency differences between early and late emerging flies were highly correlated between the host races (r = 0.54, p < 0.0001 for all 10,241 SNPs genotyped; r = 0.66, p < 0.0001 for 4244 of these SNPs mapped to chromosomes 1–5; Ragland et al., 2017). Therefore, we report results using averaged allele frequencies between the host races to estimate the mean allele frequency differences between emergence bulks (Ragland et al., 2017).

#### 2.3.3 Pre-winter selection experiments

The RAD-seq data from the pre-winter selection experiments in the hawthorn and apple races from Egan et al. (2015) and Doellman et al. (2019), respectively, were used to test whether loci associated with diapause phenotypes also responded to laboratory selection. Both studies genetically compared flies surviving different pre-winter treatments in the laboratory. Briefly, pupae were reared from field-collected fruit in Grant, Michigan and exposed to either a short (7-day) or long (32-day) pre-winter warming treatment (26°C, 15:9 L:D) starting on the day of pupariation. These treatments mimic conditions more typically experienced by the hawthorn race (short pre-winter) and apple race (long pre-winter) in the field. Following the pre-winter treatment, pupae were overwintered for 30 weeks, then held at 21°C, 14:10 L:D until adult emergence. Random samples of successfully emerging flies from each treatment were sequenced.

### 2.4 Diapause intensity phenotyping

We used two methods in the experiments to assign a diapause intensity ‘class’ to an individual fly, the first of which is based on adult emergence patterns when pupae are exposed to warm temperatures without chilling (Dambroski and Feder 2007). This approach is based on an observed, bi-modal distribution of early and late emergence timing modes corresponding to non-diapause (ND) and shallow-diapause (SD) phenotypes, respectively. Individuals that did not emerge were classified as exhibiting the diapause (DIA) phenotype (the DIA phenotype has previously been referred to as chill-dependent [CD] in *R. pomonella* [Dambroski & Feder, 2007]). The second method is based on metabolic rate measured via stop-flow respirometry, which serves as a biomarker for diapause status (Ragland et al., 2009). Briefly, individual pupae were sealed with 3-ml CO_2_-free air in syringes, and after 24 hours the full 3-ml was injected into a Sable Systems (Las Vegas, Nevada, USA) flow-through respirometry system including a LI-COR (Lincoln, Nebraska, USA) LI 7000 CO_2_ sensor. Standard bolus integration calculations accounting for CO_2_ levels in empty, control syringes were performed to estimate metabolic rate in units μl CO_2_(mg/h)^−1^. Sets of pupae were sealed and measured in blocks of 30 syringes (plus 4 controls). Replicate measures were taken over a time series (details of sampling intervals below) to characterize trajectories that fit the three discrete shapes depicted in Fig. 2a,b. Most trajectories (~85–95%; Fig. S1) could clearly be assigned to one of the three shapes/phenotypes; we discarded data from individuals with ambiguous trajectories.

**Figure 2:**
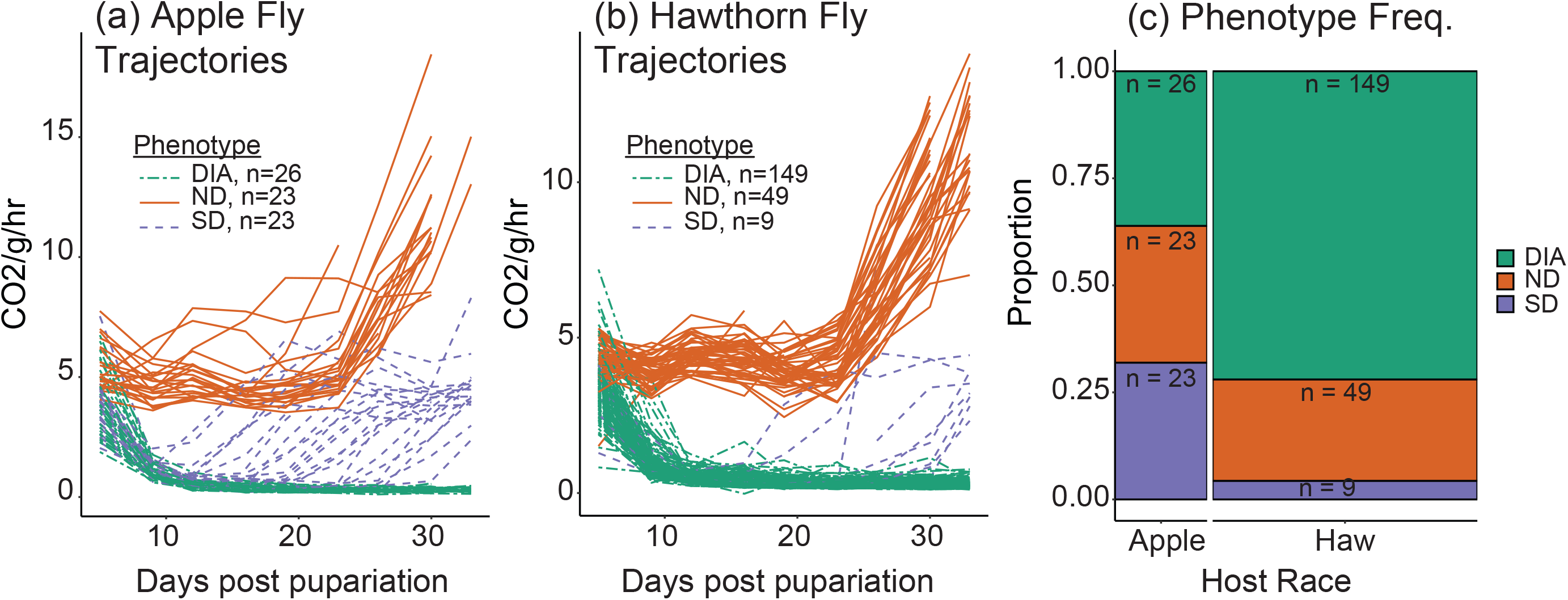
Metabolic rate (measured as CO2 production per wet mass per hour) trajectories of (a) apple-infesting host race and (b) hawthorn-infesting host race individuals exposed to simulated laboratory pre-winter conditions following pupariation. Pupariation is defined as the formation of the puparium, a hardened outer cuticle that covers the pupa during diapause and post-diapause development. Trajectories were classed as diapausing (DIA), non-diapausing (ND), and shallow diapausing (SD) based on their diagnostic trajectories. (c) Comparisons of the frequency of each diapause class within each host race, including sample sizes (n).

#### 2.4.1 Host race comparison

Pupae sampled in 2008 from field-collected apple and hawthorn fruits at Grant, Michigan were phenotyped using the metabolic rate method to compare the proportion of diapause intensity phenotypes between host races. The experiment was originally designed to provide a host race comparison and adequate sample sizes for a genetic association study in the hawthorn race. Thus, we measured three times more pupae in the hawthorn relative to the apple race. However, pupae were overwintered following measurements, and mortality during winter limited sample sizes, particularly for the rare shallow diapausing phenotype. Here, data from this experiment were used solely to compare phenotypic proportions among host races. We measured three blocks of 30 apple race pupae (90 total), and ten blocks of 30 hawthorn race pupae (300 total) twice weekly from five to 33 days post-pupariation. After accounting for mortality (some caused by handling), ambiguous trajectories, and parasitism, total sample sizes were 72 and 207 for the apple and hawthorn race, respectively (Table S1).

#### 2.4.2 Genetic associations

Pupae collected in 2009 from hawthorn fruits from Grant, Michigan were phenotyped for diapause intensity as part of a genetic association analysis. To ensure adequate sample sizes for the rarer phenotypes, two parallel experiments were performed, one based on metabolic phenotyping using stop-flow respirometry conducted at the University of Florida, and one based on adult emergence conducted at the University of Notre Dame. At the University of Florida, we used stop-flow respirometry to quantify the metabolic rate of 450 pupae (15 blocks of 30 pupae), measuring each pupa once per week from five to 75 days post pupariation, or until adult fly emergence. We excluded pupae that had been parasitized, then froze all samples including the emerged adult flies. After accounting for mortality (n = 55), parasitism (n = 149; parasitism rate was relatively high at Grant in 2009), and ambiguous metabolic rate trajectories (n = 5), we retained phenotypes for 197 total pupae or emerged adult flies (Table S2, Fig. S1).

At the University of Notre Dame, freshly pupariated larvae were held in moist vermiculite in petri dishes at 24°C and were observed three times per week for emerging flies (frozen on the day of emergence), from days five to 90 post-pupariation. The frequency distribution of emergence times was then used to determine the diapause intensity phenotype (ND, SD, or DIA). Non-diapause flies typically emerge at about 30 days post pupariation (Dambroski & Feder, 2007; Ragland et al., 2009), and the frequency distribution of emergence times showed a clear peak at around 30 days, followed by a pronounced drop-off and a broad, second mode (Fig. S2) similar to that observed by Dambroski and Feder (2007). To minimize errors in classification for the genetic association study, we only sampled ND phenotypes emerging at or earlier than 31 days, and shallow-diapause phenotypes emerging at or later than 50 days. At day 90, remaining pupae were frozen, then later the top of the puparium was removed to identify parasitized individuals that were removed from analysis. Fly pupae showing no visual markers of development were considered to be in diapause (Ragland et al., 2009).

Genotyping of phenotyped flies was carried out following the RAD-seq approach described above. We genotyped a total of 64 individuals for each diapause class phenotyped in this study (ND, SD, and DIA). Of these individuals, 37 ND, 31 SD, and 37 DIA flies were metabolically phenotyped in the experiments conducted at the University of Florida and the remaining individuals were phenotyped based on emergence timing as scored in the experiment conducted at the University of Notre Dame. Sequencing was performed at the Beijing Genomics Institute Americas on the Illumina HiSeq 2000 platform (2 full lanes), generating > 317 million 100 bp single end reads. Following data processing, we identified a common set of 7,265 variable SNPs passing quality filters that were also present in the previously published data sets on geographic variation, GWA for diapause termination, and response to laboratory selection (Doellman et al., 2019, 2018; Egan et al., 2015; Ragland et al., 2017). Quality filters included a minimum of 400x total coverage depth (across all samples), Minimum Alternate Allele (MAF) frequency > 0.05, Phred-scaled Genotype Quality (GQ) score > 20, and a non-significant test for deviations from equal representation of alleles in heterozygotes (Egan et al., 2015).

The analyses presented below were performed on these 7,265 SNPs, 3,171 of which have been mapped to the five major chromosomes of the *R. pomonella* genome (Egan et al., 2015; Ragland et al., 2017). Genotypes were polarized such that the reference allele was the allele most common in the hawthorn race at Grant, Michigan. Analyses involving loci mapped to chromosome 5 were conducted with only female individuals in each diapause class. We used this sampling design because the sex ratios in the DIA and ND groups were skewed towards females and there is evidence that chromosome 5 harbors sex determining genetic elements (M.M. Doellman & J.L. Feder, personal communication). Thus, using only a single sex in the analysis reduces the confounding effect of sex-based genetic differences.

### 2.5 Data analysis

#### 2.5.1 Linkage disequilibrium and inversion polymorphism

*Rhagoletis pomonella* has a highly structured genome that complicates genetic analysis. In particular, previous studies present evidence for multiple inversion polymorphisms on all of the five major chromosomes of the *R. pomonella* genome that produce distinct blocks of loci in high LD (Egan et al., 2015; Feder, Roethele, Filchak, Niedbalski, & Romero-Severson, 2003; Michel et al., 2010; Ragland et al., 2017). In addition to the locus-by-locus analyses described below, we also separately analyzed three different classes of linked SNPs on each chromosome categorized as displaying high (r^2^ ≥ 0.6 with one another), intermediate (0.15 ≤ r^2^ < 0.6), or low (r^2^ < 0.15 with all other SNPs) levels of composite LD identified in Ragland et al. (2017; Table S3). Chromosomes 1, 3, 4, and 5 each have a single high LD group of loci, while chromosome 2 contains eight distinct high LD clusters.

#### 2.5.2 Genetic architecture of diapause intensity

To determine what proportion of the phenotypic variation in diapause intensity is explained by heritable genetic variation, and how much of that genetic variation is accounted for by loci of major effect, we ran a Bayesian Sparse Linear Mixed Model (BSLMM) as implemented in the GEMMA program (Zhou, Carbonetto, & Stephens, 2013; more details in Supporting information 1). Bayesian model fitting allows the estimation of hyperparameters including the Proportion of the Variance in the phenotype Explained by the genotype (PVE) and the Proportion of the Genetic Variance explained by the ‘measurable’ Genetic effects (PGE). Because diapause intensity classes were categorical, we initially ran the BSLMM on pairwise comparisons of diapause classes, SD vs. DIA, SD vs. ND, and DIA vs. ND (the phenotype set equal to one or zero in the linear model) to determine how much genetic variance accounted for variation between phenotypes. These pairwise comparisons suggested that the ND and DIA classes were essentially indistinguishable; a subsequent model thus compared the pool of ND and DIA class individuals to the SD class (see Results section below). For each pairwise comparison, we also calculated a polygenic score, the mean proportion of an individual fly’s genome that contains the allele more common in flies with one of the two genotypes (Egan et al., 2015).

#### 2.5.3 Mean allele frequencies for geographic cline analysis

Diapause intensity and diapause termination are both traits that respond to natural selection imposed by seasonality (Feder, Roethele, et al., 1997; Feder, Stolz, et al., 1997). Given that seasonality also shifts with latitude (Dambroski & Feder, 2007; Feder & Bush, 1989), we also expected to see changes in allele frequencies across the four sympatric sites examined in this study. To measure geographic shifts in allele frequencies, we calculated the mean allele frequencies across loci that were significantly associated with either diapause intensity or diapause termination for each host race at all sympatric sites (Supporting information 2).

#### 2.5.4 Tests for allele frequency differences

The magnitude of allele frequency differences among groups of genotypes is proportional to the strength of genetic association (comparison among phenotype bulks) or population differentiation (comparison between random population samples). We estimated allele frequency differences between groups from genotype probabilities following methods described in Egan et al. (2015), including a non-parametric, permutation-based analysis testing whether allele frequency differences were greater (absolute value) than expected by chance (nominal alpha = 0.05; Supporting information 3). Briefly, point estimates of allele frequency differences between two groups were compared to a null distribution generated by randomly assigning group membership of each individual, thus preserving LD relationships among loci.

We used a similar permutation approach to test whether the number of SNPs with nominally significant allele frequency differences on a particular chromosome-LD was greater than expected by chance alone (Ragland et al. 2017; Supporting information 3). For this test, we calculated the percentage of SNPs within each chromosome-LD group that were significantly associated with a phenotype and compared that value against a null distribution of percentages generated using a permutation-based method (Supporting information 3).

#### 2.5.5 Correlations of allele frequency differences

We estimated correlation relationships between various allele frequency difference estimates to assess whether loci associated with diapause intensity or diapause termination also associated with previously observed genetic differences between host races, across geography, and/or pre and post-laboratory selection. These analyses underlie the triangulation approach of combining evidence for the role of sets of loci in the process of adaptation and host race differentiation, and include all combinations detailed in Table 2. We used a permutation approach similar to the one described above, comparing the empirical point estimate of Spearman’s rank correlation between allele frequency differences calculated from two different experiments against a null distribution of Spearman rank correlations estimates derived from permuted whole fly genotypes (Doellman et al., 2019, 2018; Supporting information 3).

**Table 2:**
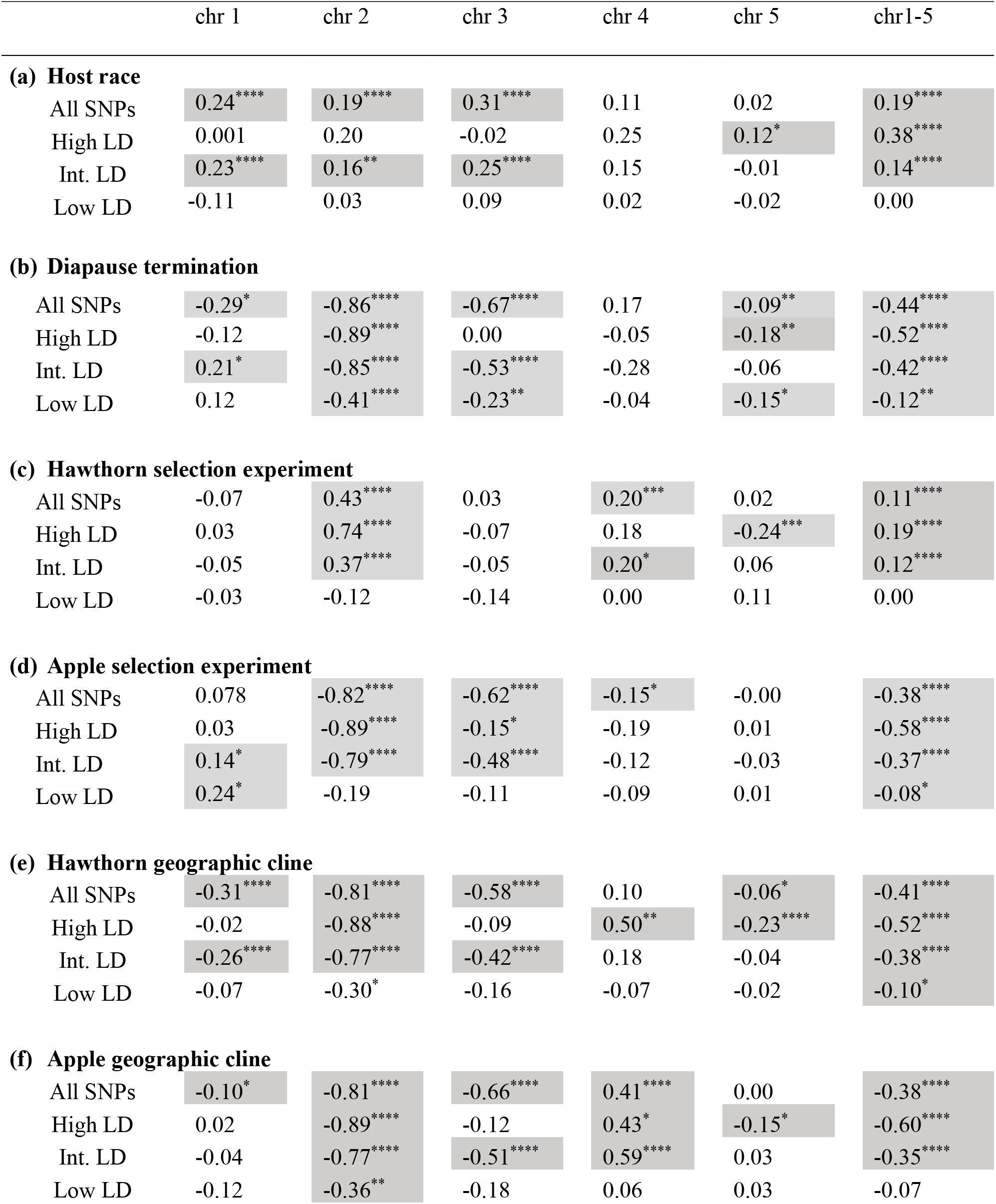
Correlation coefficients (r) of the strength of the associations between diapause intensity (allele frequency differences between SD and the combined DIA and ND groups) and allele frequency differences between various groups quantifying A) host race divergence (apple race – hawthorn race), B) strength of the association with diapause termination (early – late emergence bulks), strength of the response to selection on C) the hawthorn race and D) the apple race (32 day treatment – 7 day treatment for both) sampled from Grant, MI, and the strength of geographic divergence for E) the apple race and F) the haw race (Grant, MI – Urbana, IL for both). Results are given for all SNPs, and the High, Intermediate (Int.), and Low LD classes of loci for each chromosome considered separately, as well as for the five together (chr 1-5). * = P ≤ 0.01; ** = P ≤ 0.01; *** = P ≤ 0.001; **** ^=^ P ≤ 0.0001; significant relationships are highlighted in grey shaded boxes, as determined by Monte Carlo simulations. See Table S3 for number of SNPs genotyped in a class.

|  | chr 1 | chr 2 | chr 3 | chr 4 | chr 5 | chr1-5 |
| --- | --- | --- | --- | --- | --- | --- |
| **(a) Host race** |  |  |  |  |  |  |
| All SNPs | 0.24**** | 0.19**** | 0.31**** | 0.11 | 0.02 | 0.19**** |
| High LD | 0.001 | 0.20 | -0.02 | 0.25 | 0.12* | 0.38**** |
| Int. LD | 0.23**** | 0.16** | 0.25**** | 0.15 | -0.01 | 0.14**** |
| Low LD | -0.11 | 0.03 | 0.09 | 0.02 | -0.02 | 0.00 |
| **(b) Diapause termination** |  |  |  |  |  |  |
| All SNPs | -0.29* | -0.86**** | -0.67**** | 0.17 | -0.09** | -0.44**** |
| High LD | -0.12 | -0.89**** | 0.00 | -0.05 | -0.18** | -0.52**** |
| Int. LD | 0.21* | -0.85**** | -0.53**** | -0.28 | -0.06 | -0.42**** |
| Low LD | 0.12 | -0.41**** | -0.23** | -0.04 | -0.15* | -0.12** |
| **(c) Hawthorn selection experiment** |  |  |  |  |  |  |
| All SNPs | -0.07 | 0.43**** | 0.03 | 0.20*** | 0.02 | 0.11**** |
| High LD | 0.03 | 0.74**** | -0.07 | 0.18 | -0.24*** | 0.19**** |
| Int. LD | -0.05 | 0.37**** | -0.05 | 0.20* | 0.06 | 0.12**** |
| Low LD | -0.03 | -0.12 | -0.14 | 0.00 | 0.11 | 0.00 |
| **(d) Apple selection experiment** |  |  |  |  |  |  |
| All SNPs | 0.078 | -0.82**** | -0.62**** | -0.15* | -0.00 | -0.38**** |
| High LD | 0.03 | -0.89**** | -0.15* | -0.19 | 0.01 | -0.58**** |
| Int. LD | 0.14* | -0.79**** | -0.48**** | -0.12 | -0.03 | -0.37**** |
| Low LD | 0.24* | -0.19 | -0.11 | -0.09 | 0.01 | -0.08* |
| **(e) Hawthorn geographic cline** |  |  |  |  |  |  |
| All SNPs | -0.31**** | -0.81**** | -0.58**** | 0.10 | -0.06* | -0.41**** |
| High LD | -0.02 | -0.88**** | -0.09 | 0.50** | -0.23**** | -0.52**** |
| Int. LD | -0.26**** | -0.77**** | -0.42**** | 0.18 | -0.04 | -0.38**** |
| Low LD | -0.07 | -0.30* | -0.16 | -0.07 | -0.02 | -0.10* |
| **(f) Apple geographic cline** |  |  |  |  |  |  |
| All SNPs | -0.10* | -0.81**** | -0.66**** | 0.41**** | 0.00 | -0.38**** |
| High LD | 0.02 | -0.89**** | -0.12 | 0.43* | -0.15* | -0.60**** |
| Int. LD | -0.04 | -0.77**** | -0.51**** | 0.59**** | 0.03 | -0.35**** |
| Low LD | -0.12 | -0.36** | -0.18 | 0.06 | 0.03 | -0.07 |

#### 2.5.6 X-fold tests of parallel allelic variation between GWA studies

If sets of loci are strongly differentiated in two comparisons but at relatively uniform magnitudes (e.g., associated with diapause termination and initiation), the above correlation analysis may fail to detect a relationship. To account for this scenario, we also implemented the x-fold test as described in Chaturvedi et al. (2018; see our Supporting information 4). This approach tests whether more loci than would be expected by chance exhibit the same or opposite sign allele frequency differences for two comparisons (here, differences between emergence bulks and differences between diapause intensity classes), using permuted whole fly genotypes to generate the null distribution. Significant values falling in the lower/upper 2.5/97.5% quantiles of the null distribution suggest that loci within the set tend to be associated with, in this case, both diapause traits (termination and intensity).

## 3. RESULTS

### 3.1 Variation in diapause intensity phenotypes between host races

Three discreet diapause intensity phenotypes were apparent from metabolic rate trajectories, and these differed in frequency between the apple and hawthorn races collected from Grant, Michigan (Fig. 2). As observed in Ragland et al. (2009), diapausing pupae (DIA) displayed a rapid decrease in metabolic rate post-pupariation reaching a stable baseline in under 10 days (Fig. 2a,b, green dot-dashed lines), whereas non-diapause pupae (ND) never dropped to the low diapause baselines, and after about 20 days, they increased in metabolic rate as adult development began (Fig. 2a,b, orange solid lines). We also found a third, shallow diapausing (SD) class that dropped to a low diapause baseline initiating diapause, and approximately 10 – 30 days thereafter increased at a metabolic rate consistent with diapause termination (Fig. 2a,b, purple dashed lines). This latter, shallow diapause phenotype occurred at a markedly, and significantly higher proportion in the apple (32%) compared to the hawthorn race (4%) at Grant, MI (Fig. 2c.; *χ*^2^ = 47.85, df = 2, *p* < 0.0001).

### 3.2 Genetic associations with the SD diapause intensity phenotype

Comparing the genetic variance explained (PVE) by BSLMMs contrasting all pairwise combinations of the DIA, SD, and ND phenotypes suggested that the group of flies with the SD phenotype was genetically the most distinct, while DIA and ND groups were genetically most similar. The 95% equal tailed probability interval (ETPI) for estimates of percent variance in the phenotype explained by variation in all SNPs (PVE) for the SD vs. DIA and SD vs. ND comparisons were 0.53–0.99, centered on 0.87, and 0.57–0.99, centered on 0.88, respectively. The 95% ETPI for estimates of PVE in the DIA vs. ND comparison was 0.005–0.85, centered on 0.24 (Fig. 3a). In contrast, all estimates of the proportion of variance explained by loci of measurable effect (PGE) overlapped zero, providing no evidence for loci of large effect on any phenotypic combination (Table S4).

**Figure 3:**
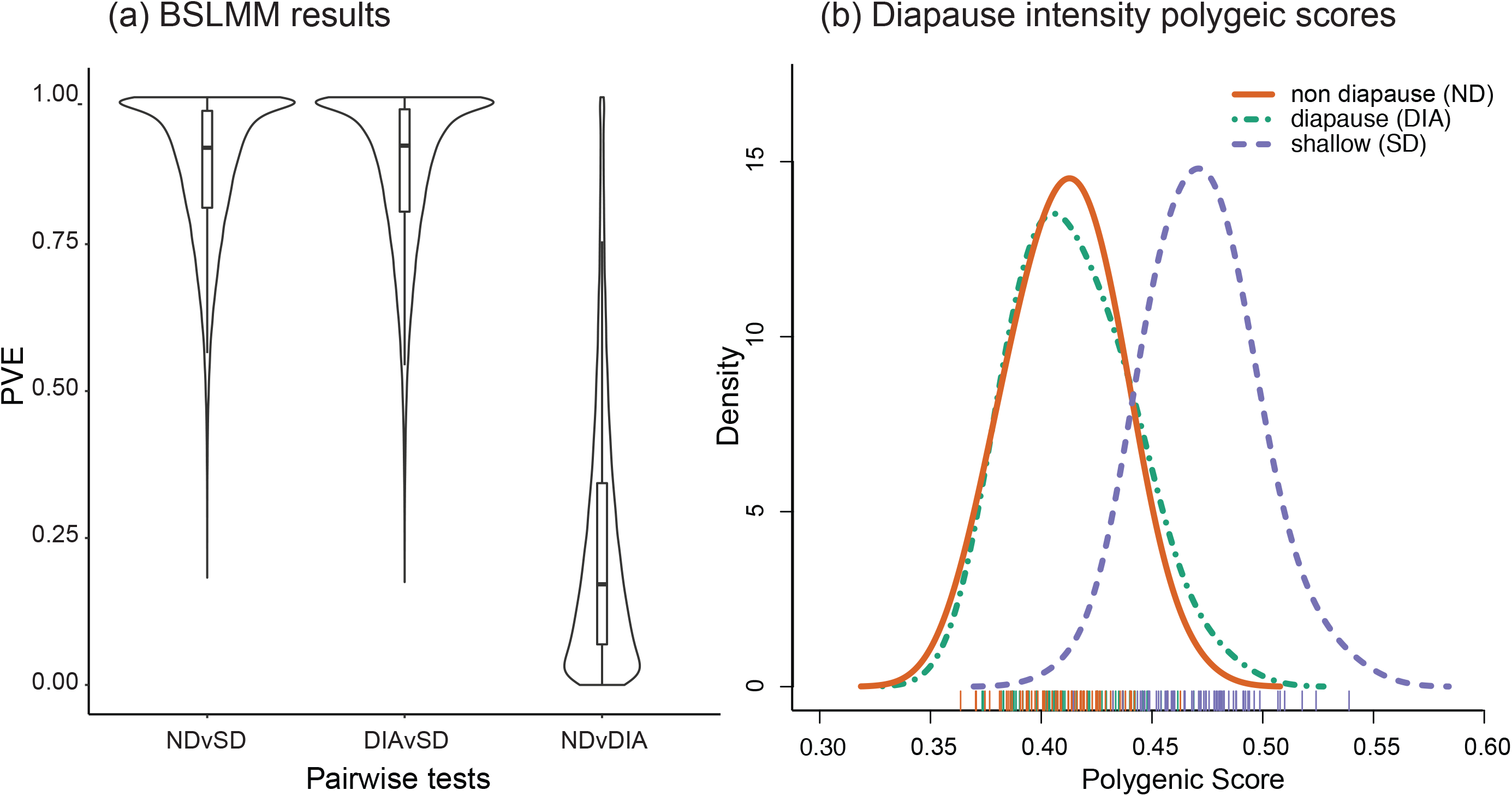
Genetic differences between diapause intensity classes SD (n=64), DIA (n=64), and ND (n=64). The left panel (a) is a violin plot of the posterior probability distribution of proportion of phenotypic variance explained by the total genetic variance (PVE) for three BSLMMs incorporating data for only two phenotypes at a time: 1) ND vs. SD, 2) DIA vs. SD, and 3) ND. vs. DIA, including boxplots of the first, second, and third quartiles surrounded by a kernel density plot. The right panel (b) displays kernel density plots of the distributions of individual polygenic scores (mean proportion of SD-like alleles that an individual has in their genome) incorporating only loci that had posterior inclusion probability (PIP) above 0.01% in at least one of the three BSLMMs.

Estimating the distribution of polygenic scores, the mean proportion of an individual fly’s genome that contains the allele more common in flies expressing the SD phenotype (Egan et al. 2015), also supported the group of flies with the SD phenotype as genetically distinct from DIA and ND groups. Only loci that had a posterior inclusion probability (PIP) above 0.01% in at least one of the three pairwise comparison BSLMMs (n = 2,220) were used to calculate the polygenic scores, whose distributions for each phenotype are illustrated in Figure 3b. Consistent with the results of the pairwise BSLMM models described above, the distribution of polygenic scores for the SD phenotype was shifted towards higher values (Komolgorov Smirnov tests - SD vs. DIA: *D* = 0.83, *p* < 0.0001. SD vs. ND: *D* = 0.92, *p* < 0.0001), while distributions for the DIA and ND phenotypes were largely overlapping (Komolgorov Smirnov test - DIA vs. ND: *D* = 0.10, *p* = 0.86)

The combined evidence from the BSLMM and polygenic score distributions suggested that most genetic variance was associated with the distinction between the group of flies with the SD phenotype as compared to the groups of flies with either the DIA or ND phenotypes, which were relatively genetically homogenous. Thus, all subsequent analyses of genetic associations compared SD phenotypes to a combined group of DIA and ND phenotypes (DIA+ND).

### 3.3 Chromosomal distribution of diapause intensity-associated SNPs

Significant allele frequency differences between the SD and DIA + ND groups, evidence for genetic association with diapause intensity, were largely confined to chromosomes 2 and 3 Figure 4). Significant percentages of SNPs in the high LD clusters on chromosomes 2 (81%) and 3 (55%) exhibited significant allele frequency differences (Fig. 4a). Significant percentages of SNPs significantly associated with diapause intensity were also observed within the intermediate and low LD groups on chromosome 2 (65% and 18%, respectively), the intermediate LD group on chromosome 3 (17%), and the high LD group on chromosome 5 (9%; Fig. 4a). There was no significant excess of SNPs significantly associated with diapause intensity anywhere on chromosome 1 (low LD: 7%, Int. LD: 4%, high LD: 0%), the low LD region on chromosome 3 (3%), anywhere on chromosome 4 (low LD: 5%, int. LD: 3%, high LD: 3%) and the intermediate and low LD regions on chromosome 5 (low LD: 8%, int. LD: 6%; Fig. 4a).

**Figure 4:**
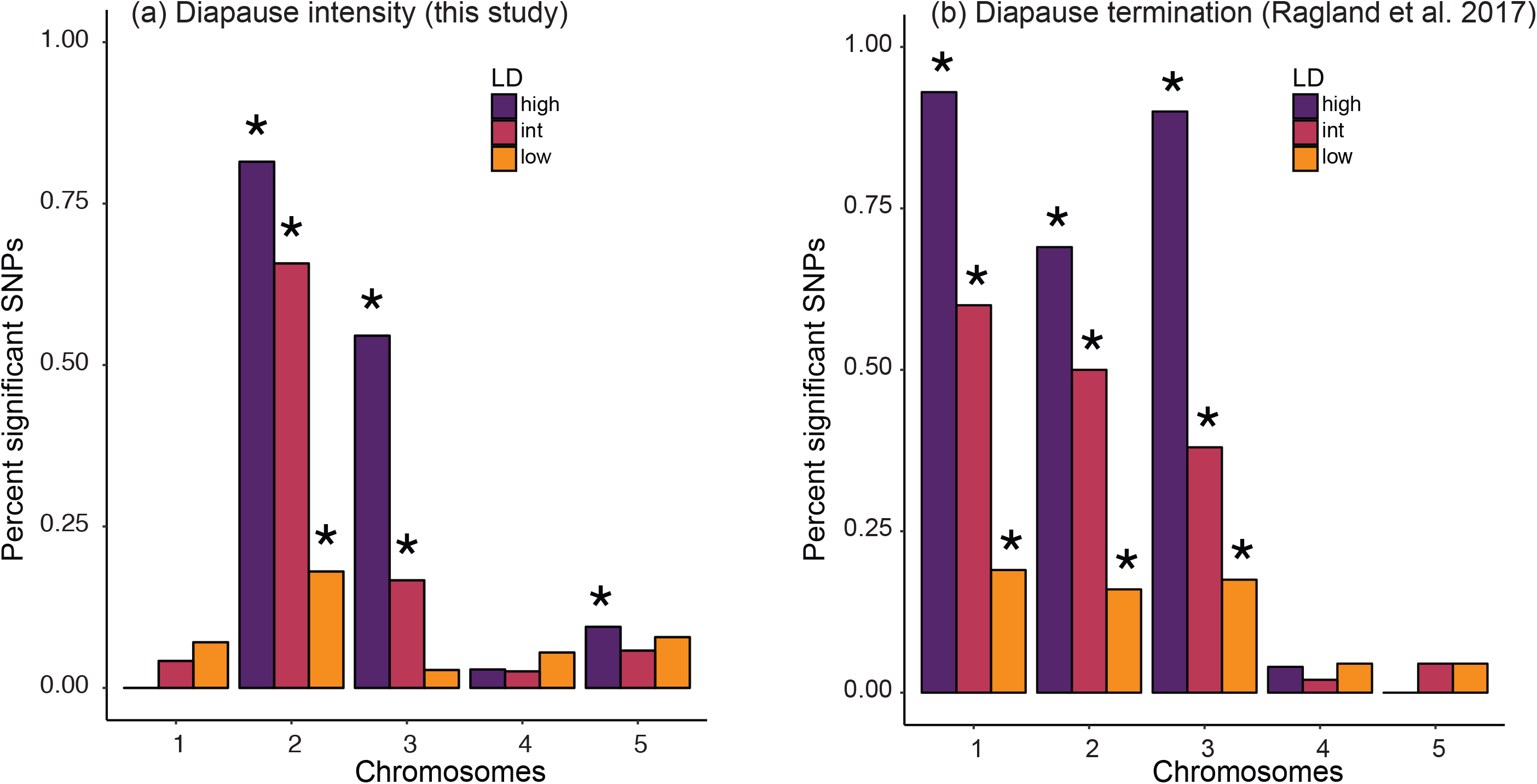
Percentage of SNPs significantly associated with diapause intensity (a) and diapause termination (b) in each LD region on each chromosome. The diapause intensity data was genotyped in this study (n=64 for each diapause class), while the diapause termination data was genotyped in Ragland et al. (2017; n=48 for each host race and diapause termination group [early vs. late]). For the diapause intensity and diapause termination GWAS the statistical significance of SNPs was determined through non-parametric permutation tests (Supporting information 3). A star indicates a significant (*p* < 0.05) excess of significantly associated SNPs compared to null expectations.

Ragland et al. (2017) reported an excess of SNPs associated with diapause termination on chromosomes 1, 2 and 3. Although these SNPs were also largely located within high LD regions, significant excesses of SNPs associated with diapause termination were also observed for the intermediate and low LD regions on chromosomes 1-3 (Fig. 4b). Thus, loci on chromosomes 2 and 3 were associated with both diapause intensity and diapause termination (particularly in the high LD regions), whereas most loci on chromosome 1 were associated with diapause termination, but not with diapause intensity.

### 3.4 Host race divergence for diapause-associated loci

Genetic variation associated with both diapause phenotypes was also significantly related to host race divergence at Grant, MI, though in a chromosome-dependent manner. Allele frequency differences between the SD and DIA + ND diapause classes were significantly positively associated with allele frequency differences between host races (Table 2a). Here, a positive relationship indicates a genetic association between the SD phenotype and the apple race (Fig. 5a-c). The variation underlying this relationship was largely confined to chromosomes 1 through 3 (Fig. 5a-c; Supporting information Fig. S3 for chromosomes 4 and 5). For each of those chromosomes, only the intermediate LD regions, when considered alone, exhibited a significant relationship between diapause intensity and host race (Table 2a). In contrast, the results of Ragland et al. (2017) suggest a negative correlation between associations with diapauses termination and with host race (Table 3). This relationship also varied across chromosomes, with the strongest relationships between phenotypic- and host-associations present on chromosomes 1 and 3 (Table 3).

**Table 3:** Correlation coefficients (r) of the strength of the associations between diapause termination (allele frequency difference between early minus late emergence bulks) and host race divergence (allele frequency differences between the apple minus hawthorn race) at Grant, MI. Results are given for all SNPs, and the High, Intermediate (Int.), and Low LD classes of loci for each chromosome considered separately, as well as for the five together (chr 1-5). * = P < 0.01; ** = P < 0.01; *** = P < 0.001; significant relationships, are highlighted in grey shaded boxes, as determined by Monte Carlo simulations.

|  | chr 1 | chr 2 | chr 3 | chr 4 | chr 5 | chr1-5 |
| --- | --- | --- | --- | --- | --- | --- |
| All SNPs | -0.63* | -0.08 | -0.46** | 0.07 | 0.02 | -0.41*** |
| High LD | -0.08 | -0.09 | -0.09 | -0.14 | 0.07 | -0.56* |
| Int. LD | -0.50*** | -0.05 | -0.40** | 0.18 | 0.03 | -0.28*** |
| Low LD | -0.27* | 0.05 | -0.06 | -0.02 | 0.03 | -0.06 |

**Figure 5:**
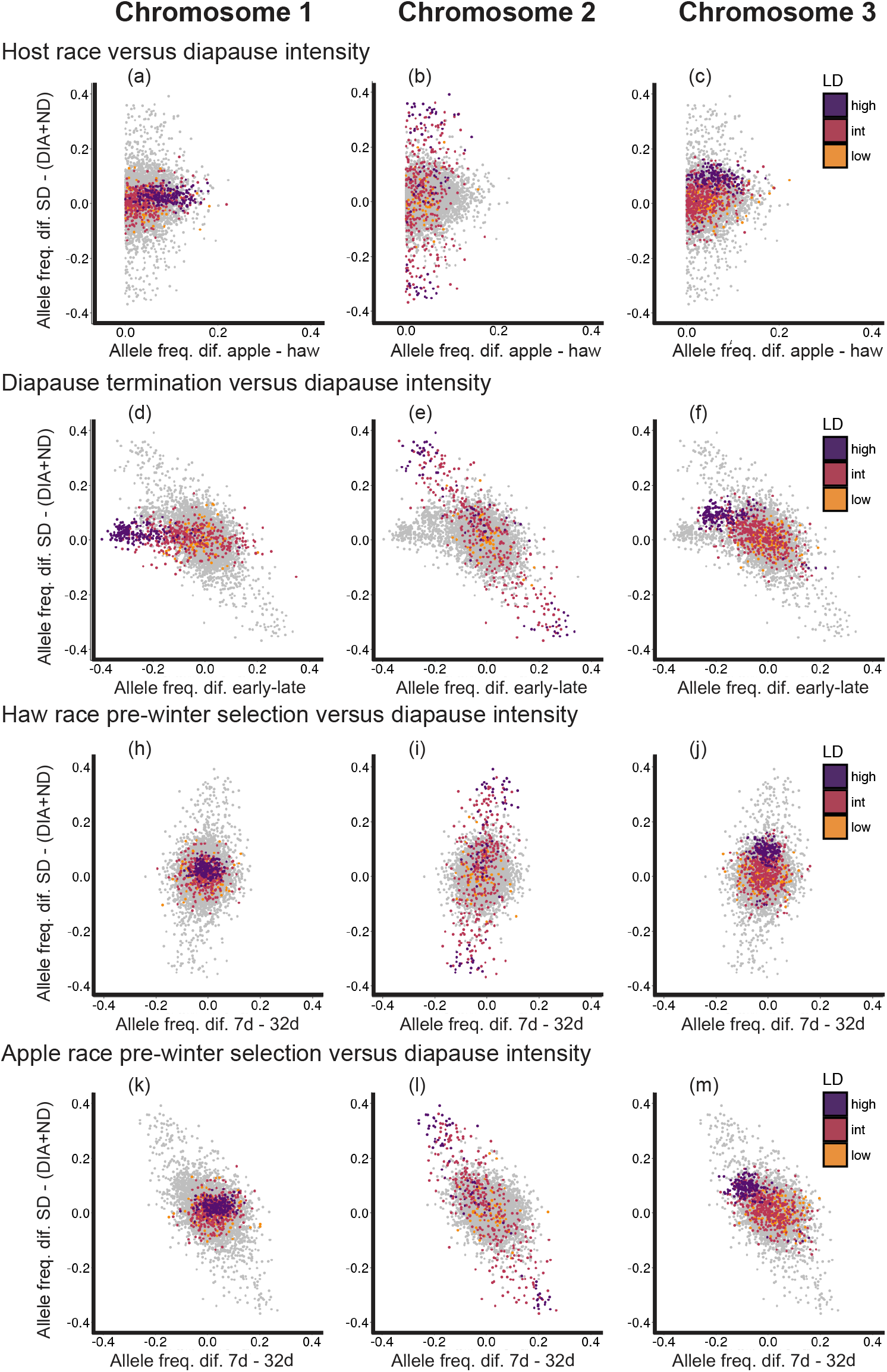
Relationships between the strength of genetic associations (magnitude of allele frequency differences) with diapause phenotypes, selection experiments, and host race differences. We tested relationships between the genetic association with diapause intensity (measured as the difference in allelic variation between the SD class and the pooled DIA and ND classes [SD − [DIA + ND]]; n = 64 for each diapause class) and host race differences (apple host race – hawthorn host race; n = 48 for the apple race, n = 54 for the haw race; a – c), diapause termination (early – late; n = 48 for both early and late groups; d – f), hawthorn fly selection experiment (7 day – 32 day; n=54 for the 7-day treatment, n = 47 for the 32 day treatment; g – i), and apple fly selection experiment (7 day – 32 day; n = 48 for the 7 day treatment, n = 41 for the 32 day treatment; j–l). Light grey dots represent all 7,265 genotyped SNPs while orange dots are low LD loci, red dots are intermediate LD loci, and purple dots are high LD loci on chromosome 1 (1st column; a,d,h,k), chromosome 2 (2nd column; b,e,i,l), and chromosome 3 (3rd column; c,f,j,m). All loci were polarized to the allele at the highest frequency in the hawthorn fly population at Grant, MI; thus, host race differences in allele frequency were all positive.

### 3.5 Genetic association between diapause intensity and termination

Genetic variation associated with diapause intensity was significantly and negatively related to the pattern of genetic variation in diapause termination across all 7,265 SNPs examined in this study (Table 2b; Fig. 5d-f). Here, a negative relationship indicates a genetic association between the SD phenotype and late diapause termination (and by extension an association between DIA + ND and early diapause termination). The overall relationship across all loci was primarily driven by variation in loci located on chromosomes 2 and 3 in high (chromosome 2) and intermediate (chromosomes 2 and 3) LD groups (Fig. 5e,f; Table 2b). x-fold enrichment tests also supported an excess of loci associated with both phenotypes on chromosomes 1, 2, 3, and 5 though primarily for loci located in high and intermediate LD groups on chromosomes 1, 2, and 3 (Fig. 6). Though the x-fold test suggested that loci associated with diapause intensity also tended to associate with diapause termination on chromosome 1, there were many more loci on chromosome 1 associated with diapause termination (n = 450) compared to just a few associated with diapause intensity (n = 21; Fig. 7a). Meanwhile, there was substantial overlap in the loci significantly associated with diapause termination and diapause intensity on chromosome 2 (n=201) and chromosome 3 (n=127; Fig 7b-c).

**Figure 6:**
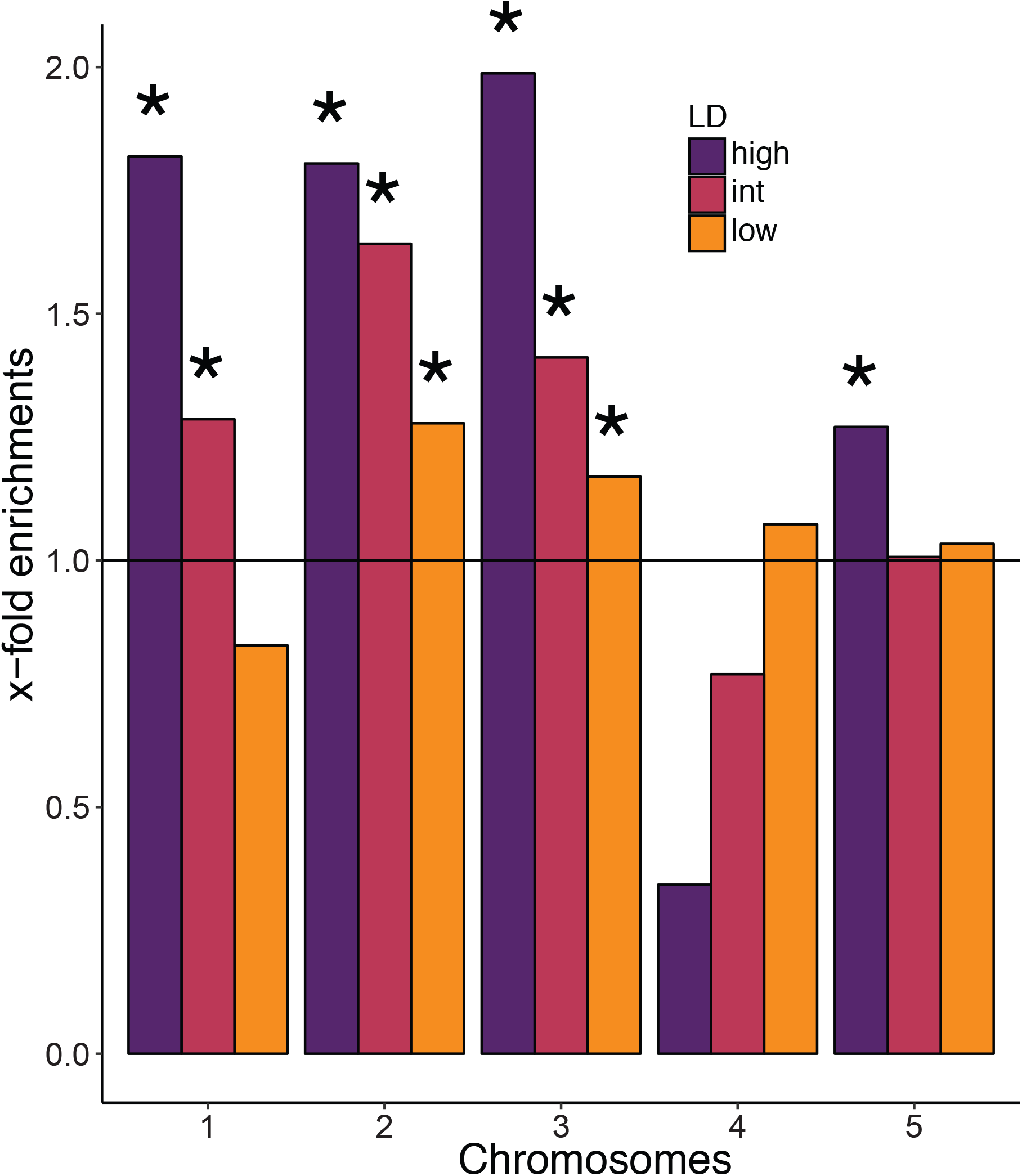
X-fold enrichments for shared signs of allele frequency differences (sign concordance) between diapause intensity associations (allele frequency differences; SD − (DIA + ND); n = 64 for each diapause class) and diapause termination associations (allele frequency differences; early – late emergence bulks; n=48 for each emergence bulk). Results are shown for each LD group (low, intermediate [int], and high) for each of the 5 chromosomes. The null expectation is shown with a solid horizontal line. Stars indicate sign concordance statistically significantly greater than expected by chance alone (p < 0.05).

**Figure 7:**
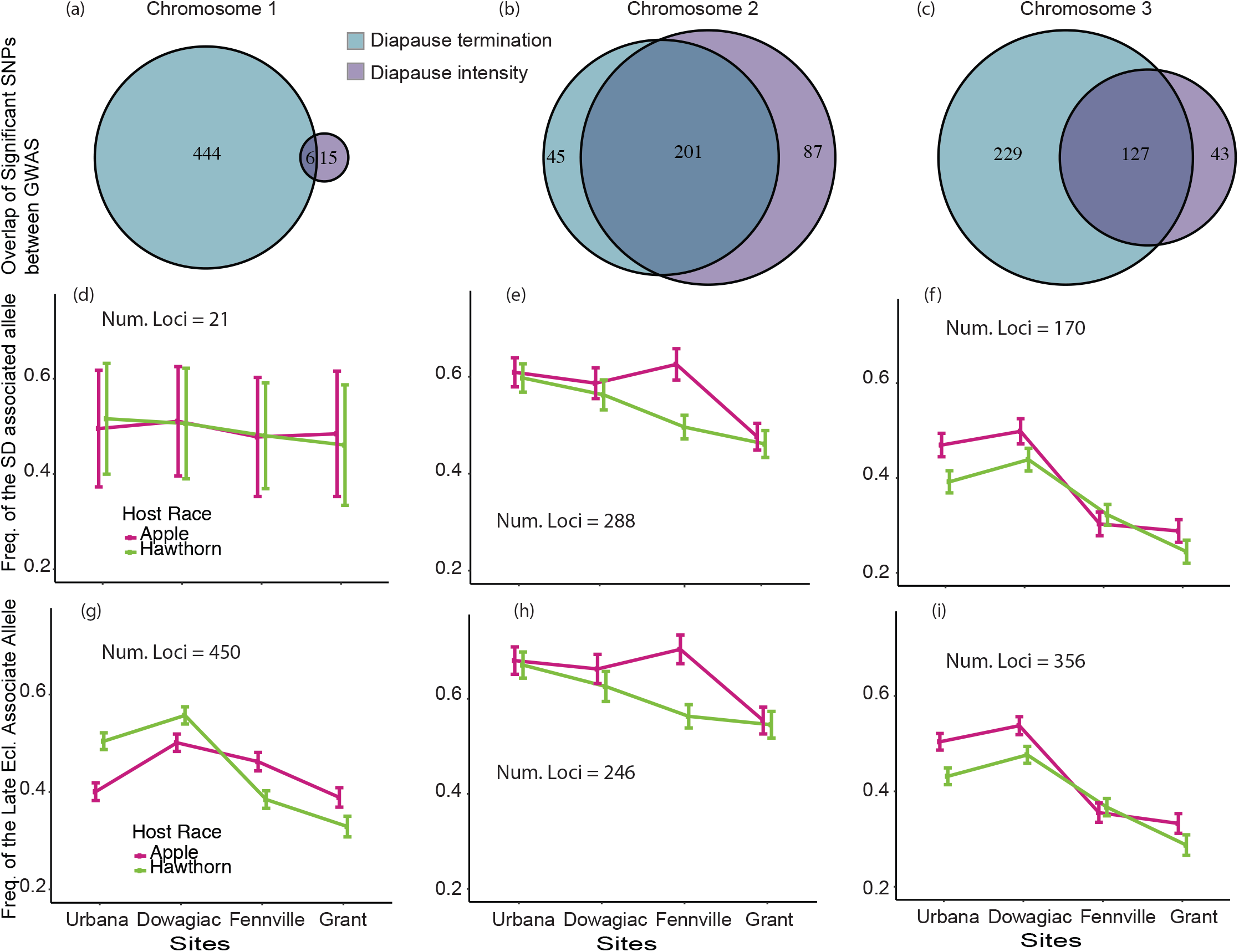
(a,b,c) Venn diagrams illustrating number of loci associated with either diapause termination (blue), diapause intensity (purple), or both (overlapping regions). (d,e,f) Mean (95% percent confidence intervals) frequencies of the allele associated with the SD phenotype at each geographic location for loci significantly associated with the diapause intensity phenotype. (g,h,i) Mean (95% confidence intervals) frequencies of the allele associated with the late diapause termination phenotype at each geographic location for loci significantly associated with the diapause termination phenotype. Results are presented for chromosomes 1 (a,d,g), 2 (b,e,h), and 3 (c,f,i), and for both hawthorn race (green lines) and apple race (pink lines) flies. The number of loci used to calculate the mean allele frequencies for each chromosome are reported on each plot.

### 3.6 Selection response for diapause-associated loci

Allele frequency differences between the SD and DIA + ND diapause classes were significantly positively related to the differences between the 7-day and 32-day treatments in the hawthorn race pre-winter selection experiment (Fig. 5h-j, Table 2c) but were significantly negatively related to allele frequency differences in the apple race selection experiment (Fig. 5k-m, Table 2d). Here, a positive relationship indicates a genetic association between the SD phenotype and apple race individuals surviving the 7-day prewinter treatment, while a negative relationship indicates the reverse. For both experiments, loci on chromosome 2 largely drove the significant correlation (Fig. 5g-m, Table 2c,d) and to a lesser extent loci on chromosome 3 (Fig. 5g-m). As with the relationship between diapause intensity and diapause termination, loci in high and intermediate LD groups were most strongly associated with both diapause intensity and the response to selection (Table 2a-d).

### 3.7 Clinal variation for diapause-associated loci

Allele frequencies for loci significantly associated with diapause intensity (SD vs. DIA + ND) on chromosome 1 did not change appreciably with geography (Fig. 7d), while frequencies of the SD-associated alleles on chromosomes 2 and 3 decreased with increasing latitude in both the hawthorn and apple races (Fig. 7e,f). Allele frequencies for loci on chromosomes 1 - 3 significantly associated with diapause termination also varied with geography, with the frequency of the allele associated with later diapause termination generally increasing with decreasing latitude (Fig. 7g-I; Doellman et al., 2019). Clines were not completely linear, with notable peaks of allele frequency at the Fennville, Michigan site. Moreover, clines were not identical between host races. In particular, separate clines for the apple and hawthorn races tended to converge with increasing latitude (termination-associated loci on chromosome 1 and 3, and intensity-associated loci on chromosome 3) such that differences between host races were most pronounced at Urbana, IL, the lowest latitude site (see below).

The clinal variation in allele frequencies is further supported by the significant negative relationship between diapause intensity and geographic differences within each of the host races between the Grant, MI and Urbana, IL sympatric sites (Table 2e,f). Here, a negative relationship indicates a genetic association between the SD phenotype and the southern sympatric site Urbana, IL. For both host races, loci on chromosomes 2 and 3. exhibited the strongest relationships between diapause intensity and geography (Table 2e,f).

## 4. DISCUSSION

### 4.1 Historic, clinal selection and antagonistic pleiotropy

We provide evidence that historical, spatially variable selection on two ecologically important traits favored LD between associated genetic loci or pleiotropy in a direction that is antagonistic to contemporary selection driving rapid adaptation in *R. pomonella*. Patterns of current genomic variation thus appear to reflect both this historical selection favoring genetic correlations and contemporary selection to break up genetic correlations. Below, we provide a summary of the combined inferences that lead to these conclusions, then discuss how these results inform our understanding of the genetic architecture of genetic correlations and adaptive divergence.

First, results of the BSLMM and locus-by-locus analyses confirm that, in addition to previously described variation in diapause termination, there is segregating variation for diapause intensity (addressing question 1). Moreover, this variation is concentrated in intermediate and high LD groups on three of the five *R. pomonella* chromosomes, and colocalizes with variation associated with diapause termination on chromosomes 2 and 3. Second, the SD phenotype occurred at a higher frequency in the apple race (Fig. 2), as did the frequencies of alleles associated with SD (addressing question 2). In addition, the observed correlations between the strength and direction of genetic associations with diapause intensity and host race differences on chromosomes 2 and 3, suggest that diapause intensity is differentially selected between the host races, with the SD phenotype favored in the apple race. Third, loci associated with diapause intensity were also largely associated with diapause termination, as evidenced by significant correlations across all SNPs (Table 2, Fig. 5d-f) and high x-fold values (Fig. 6), suggesting either pleiotropy or strong LD among the genetic elements controlling these traits (addressing question 3). Furthermore, this genetic relationship supports a phenotypic association between SD and late diapause termination (and DIA/ND with early diapause termination). While such a trait combination matches the prediction for long term historical, clinal selection (Fig. 1c) in the ancestral hawthorn race, it is antagonistic to the direction of selection in the recently formed apple race. Fourth, we observed significant correlations between the strength of the genetic association with diapause intensity and the genetic response in the pre-winter selection experiments, indicating that segregating variation for diapause intensity is likely selected upon during the pre-winter period in nature (answering question 4). Finally, we observed that allele frequencies at loci associated with diapause intensity and termination vary with latitude in directions consistent with geographically variable, correlational selection (addressing question 5). Based on phenotypic variation for both traits across geography (Dambroski & Feder, 2007; Feder et al., 1994), we hypothesized that alleles associated with more intense diapause and later diapause termination would be favored in the South, with the alternative alleles conferring less intense diapause and earlier diapause termination in the North. Our genetic results largely follow this prediction, with mean frequencies of alleles associated with the SD and late diapause termination (emergence time) phenotypes increasing with decreasing latitude, including loci on chromosomes 2 and 3 that associate with both phenotypes (Fig. 7, Table 2e,f). Collectively, the answers to these five questions indicate that strong selection on diapause intensity during apple race formation has led to an increase in allelic variants associated with later diapause termination that appears to be going in the opposite direction from that predicted based on the earlier diapause termination of the apple race.

These results contradict previous findings that diapause intensity and diapause termination are genetically modular (largely uncorrelated) in *R. pomonella* (Ragland et al., 2017). However, the prior study used survival in a pre-winter selection experiment on the hawthorn race as a proxy for diapause intensity, while here we directly measured the diapause phenotype. Genetic variation in diapause intensity was still associated with the results of pre-winter laboratory selection experiments, as allele frequencies changed at loci associated with diapause intensity when both the apple and hawthorn races were exposed to long pre-wintering treatments in the laboratory (Fig. 5, Table 2c,d). The results of the two selection experiments were not concordant, however, with the sign of the correlation with diapause intensity differing between host races exposed to the selection experiment (Doellman et al., 2018).

The results described above suggest that specific phenotypes visible to selection may depend on genetic background, and/or on other, unmeasured traits that differ in frequency among the host races. The SD phenotype is much more common in the apple race (Fig 2c; Dambroski & Feder, 2007), and the SD-associated alleles are clearly more favored in the apple race flies surviving the 32 day, long pre-winter treatment. Moreover, fitness is most likely determined by combinations of many more traits than we can measure, as indicated by a large body of empirical data and theory (Blows & Hoffmann, 2005; Orr, 1998; Saltz et al., 2017). Thus, we might expect idiosyncratic genomic responses to a relatively non-specific selection pressure (pre-winter length) applied to genetically divergent populations (Chaturvedi et al., 2018; Gompert & Messina, 2016; Linnen, 2018; Nosil et al., 2018). Despite the ambiguity of the hawthorn race selection experiments, the results are consistent with ongoing selection in the apple race favoring SD-associated alleles that also associate with early diapause termination, through pleiotropy or linkage.

### 4.2 Genomic signatures of genetic correlation

With the advent of denser genome sampling enabled by next generation sequencing, there is growing interest in uncovering patterns of genome-wide variation underlying observed genetic correlations among trait combinations (Saltz et al., 2017). Our data provide a snapshot of genetic correlations generated by historical, correlational selection. In particular, loci on chromosomes 2 and 3 in high LD regions appear to pleiotropically affect both traits in the ancestral hawthorn race in a direction consistent with correlational selection for a positive association between diapause intensity and diapause termination. Our measure of pleiotropy based on shared genetic associations could be driven by actual pleiotropic effects of a given genetic variant, or by high LD driven by close physical proximity on the chromosome. Given that both traits are related to diapause regulation, pleiotropic effects of genes acting through highly pleotropic developmental pathways such as insulin (Sim & Denlinger, 2013) and *wnt* (Meyers et al., 2016; Wodarz & Nusse, 1998) signaling are plausible. However, *R. pomonella* also has a highly structured genome with blocks of high LD that may be associated with chromosomal inversions (Feder, Roethele, et al., 2003; Ragland et al., 2017). Most of the genetic variation that appeared pleiotropic occurred in high to intermediate LD regions (Fig. 6). Past studies have suggested that ancestral inversions from disjunct populations in Mexico have spread north via the same geographically variable selection that favors more intense diapause and later diapause termination in the South (Feder, Berlocher, et al., 2003; Feder et al., 2005). Combined, this evidence strongly hints at a prominent role for reduced recombination between co-adapted variants in mediating adaptation, a genomic signature also observed in other systems (Ayala et al., 2019; Coughlan & Willis, 2019; Lowry & Willis, 2010; Noor, Grams, Bertucci, & Reiland, 2001; Wallberg, Schöning, Webster, & Hasselmann, 2017).

Despite theory that adaptation should proceed more quickly when selection acts in the direction of positive trait correlations (i.e. where maximum genetic variation is available; Schluter, 1996), empirical work shows that genetic correlations can also be rapidly degraded by selection (Beldade, Koops, & Brakefield, 2002; Conner et al., 2011). Though, the genomic signatures of this breakdown remain unclear. At sympatric sites, natural selection is expected to favor a more intense diapause in the apple compared to the hawthorn race of *R. pomonella* (Fig. 1c). From this perspective, the observed pleiotropy and/or linkage relationships on chromosomes 2 and 3 are antagonistic to the direction of selection for diapause termination in the apple race. Thus, selection imposed by infesting apple vs. hawthorn fruit would be expected to break up the antagonistic phenotypic/genetic correlation on chromosomes 2 and 3 historically favored by geographically variable selection within the hawthorn race. Furthermore, selection should also favor new interactions or combination of alleles in the apple race that combine increased diapause intensity and earlier diapause termination.

Though we could not directly assess genetic correlations in the apple race (the phenotypic associations were measured in the hawthorn race), we can make inferences based on evolved host race differences. On chromosomes 2 and 3, we observed host divergence patterns consistent with a response to selection on increased diapause intensity in the apple race, at the expense of predicted, maladaptive effects increasing diapause termination timing (Fig. 7). This implies that the LD or pleiotropic relationships remain in the apple race. In contrast to chromosomes 2 and 3, loci on chromosome 1 appear to respond to selection for diapause termination in a manner independent of selection on diapause intensity. Phenotypic evidence clearly shows that the apple race has managed to evolve earlier diapause termination (Feder et al., 1994), and the effects of non-pleiotropic (or non-linked) loci such as those on chromosome 1 may overwhelm the maladaptive effects of pleiotropic/linked loci on chromosomes 2 and 3, assuming an additive model. However, loci on chromosome 1 cannot universally account for earlier diapause termination in the apple race, as the mean frequencies of the early-associated allele are higher in the apple race at only two of the four geographic sites (clines in mean allele frequency cross in Fig. 7g; Doellman et al., 2019). The RAD loci in this study represent only a portion of the genome and are likely biased to detect associations in higher LD regions (Lowry et al., 2017). Thus, unidentified loci in more collinear regions of the genome may play an important role in resolving the genetic paradox for why the apple race emerges earlier than the hawthorn race but possess higher frequencies of SD associated alleles correlated with later diapause termination.

### 4.3 Predicting genetic variation in natural populations from laboratory experiments

There is often some uncertainty in the relationship between a phenotype that we can measure in the laboratory versus phenotypes that are expressed and experience selection in the field (Chaturvedi et al., 2018; Nosil et al., 2018; Orsini et al., 2012; Soria-Carrasco et al., 2014). Here, the SD phenotype would appear to be maladaptive because adult flies prematurely emerge as adults when exposed to long, warm pre-winter conditions, although not as quickly as ND individuals. Premature diapause termination prior to winter leads to zero fitness because those flies would emerge when few suitable hosts or mates would be present (Feder, Roethele, et al., 1997). Therefore, the SD phenotype, although not as disadvantaged as ND phenotypes, should still be maladapted compared to the DIA phenotype. However, the genetic similarity of ND and DIA flies suggest that these two phenotypes may actually be related, with DIA flies being prone to non-diapause development and the ND phenotype representing a subset of DIA flies that realized this potential. Thus, SD-associated alleles may actually have a reduced risk compared to the immediate ND phenotype and, as conditions become progressively cooler during the fall when diapause is initiated, may rarely develop prematurely before winter in nature. The clinal data also suggest that the SD phenotype is favored under warmer, more southern conditions. Further studies monitoring development in the field will be necessary to better understand the multifarious nature of selection acting on different aspects of the diapause phenotype.

### 4.4 Antagonistic genetic effects and genomic divergence

Why does the directionality of allele frequency differences between ecologically divergent populations often fail to follow predictions? There are many possibilities, including 1) selection acts on highly polygenic traits wherein many genetic combinations can produce similar phenotypes (Boyle et al., 2017; Yeaman, 2015), 2) epistatic effects, where single-locus effects depend on the genetic background (Chandler, Chari, Tack, & Dworkin, 2014; Taylor & Ehrenreich, 2015), or 3) changing LD relationships between marker and target loci (Butlin & Smadja, 2018). Though we cannot rule out these explanations, our data suggest that antagonistic genetic effects, mediated either through locus-specific pleiotropy or through tight LD, and multivariate selection also play a role. This conclusion is also well supported by an experiment in natural populations of the monkeyflower, *Mimulus guttatus,* that initially suggested a negative relationship between flower size (a proxy for seed set and fitness) and viability (Mojica and Kelly 2010). Later work found that substantial pleiotropy exists for genes controlling flower size and development rate and that strong viability selection in the form of summer drought onset led to fast development times at the expense of producing smaller, less fecund flowers (Mojica et al. 2012; Troth et al. 2018). Furthermore, seasonal variation in drought onset can lead to large temporal fluctuations in the relative fitness benefits of fast developing, small flower phenotypes, and slow developing, large flower phenotypes (Troth et al. 2018). A greater understanding of the major environmental challenges faced by diapausing *R. pomonella* pupae in the field with regard to conditions before, during, and after winter will be needed to untangle the multifarious nature of selection on the lifecycles of the apple and hawthorn host races within and across geographic locations.

## 5. SUMMARY

In this study, we have provided evidence that the consistently recorded, paradoxical pattern of the apple race of *R. pomonella* harboring increased frequencies of late diapause termination timing alleles is partially a result of a correlated response to selection for pre-winter diapause intensity. Geographic clinal variation in genetic loci associated with pre-winter diapause intensity and post-winter diapause termination supports the hypothesis that historic, geographic selection in the ancestral hawthorn race has favored trait combinations and shaped genetic variation that is antagonistic to the direction of selection for diapause termination during formation of the apple race. These findings suggest that incomplete knowledge of the multivariate trait combinations responding to selection can sometimes limit our ability to accurately predict evolution. Future work will need to carefully quantify the contribution of antagonistic pleiotropy or LD to estimates of genomic predictability and evaluate what genetic mechanisms within *R. pomonella* have allowed the apple race to evolve around this apparent genetic constraint.

## Supporting information

Supplemental ISunformation

## ACKNOWLEDGMENTS

Withheld for blind review.

## Notes

### Competing Interest Statement

The authors have declared no competing interest.

