## Supplemental Figures 1-4 for "The genomics of trait combinations and their influence on adaptive divergence"

**Supporting Figures 1-4**

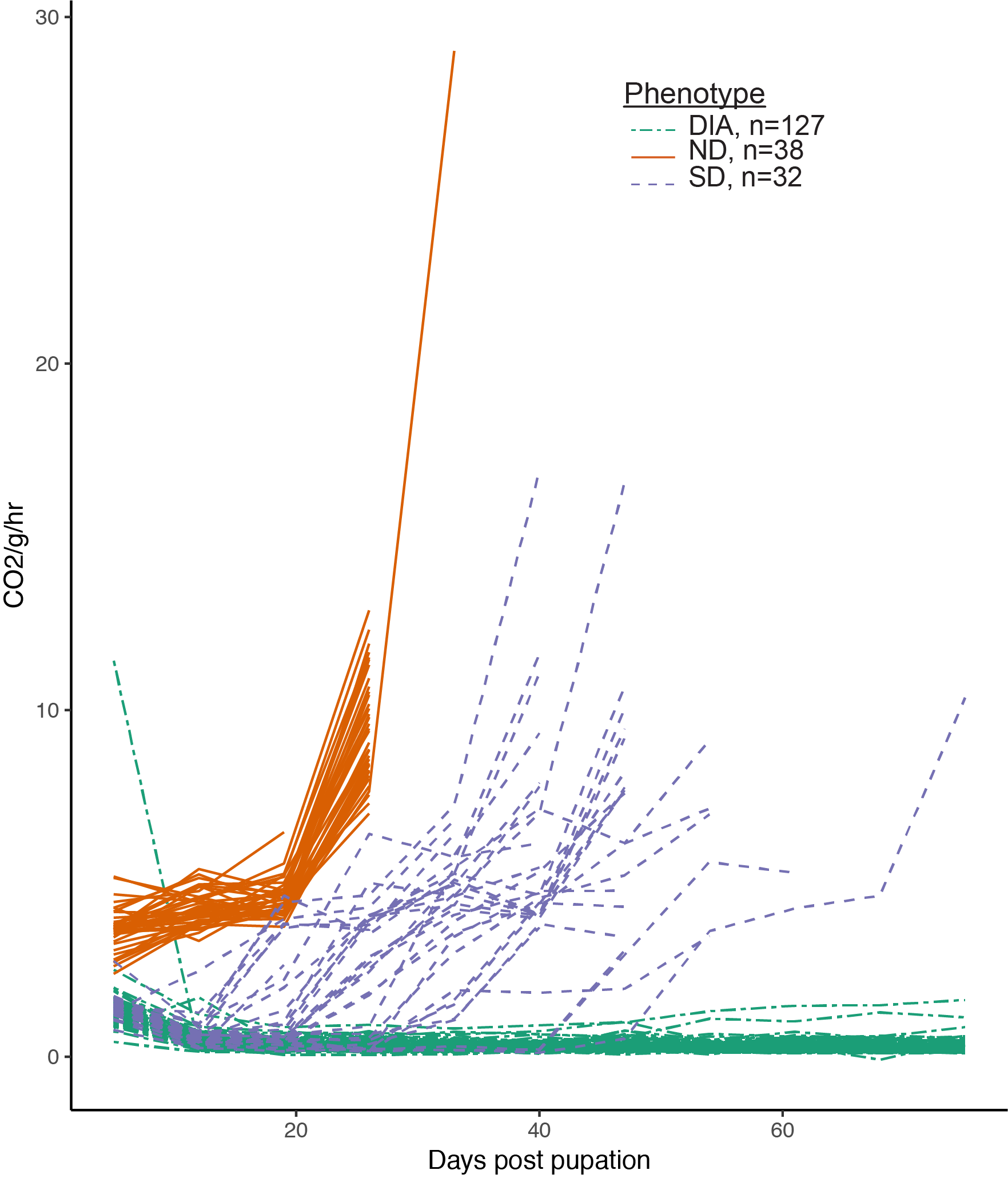

**Supporting Figure 1**: Metabolic rate trajectories of hawthorn race pupae exposed to long pre-winter conditions in the stop-flow respirometry experiment performed at the University of Florida. Data correspond with numbers reported in Table S2. Green lines indicate individuals classed as non-diapausing (ND), blue lines indicate individuals classed as shallow diapause (SD), and red lines indicate individuals classed as diapause (DIA).

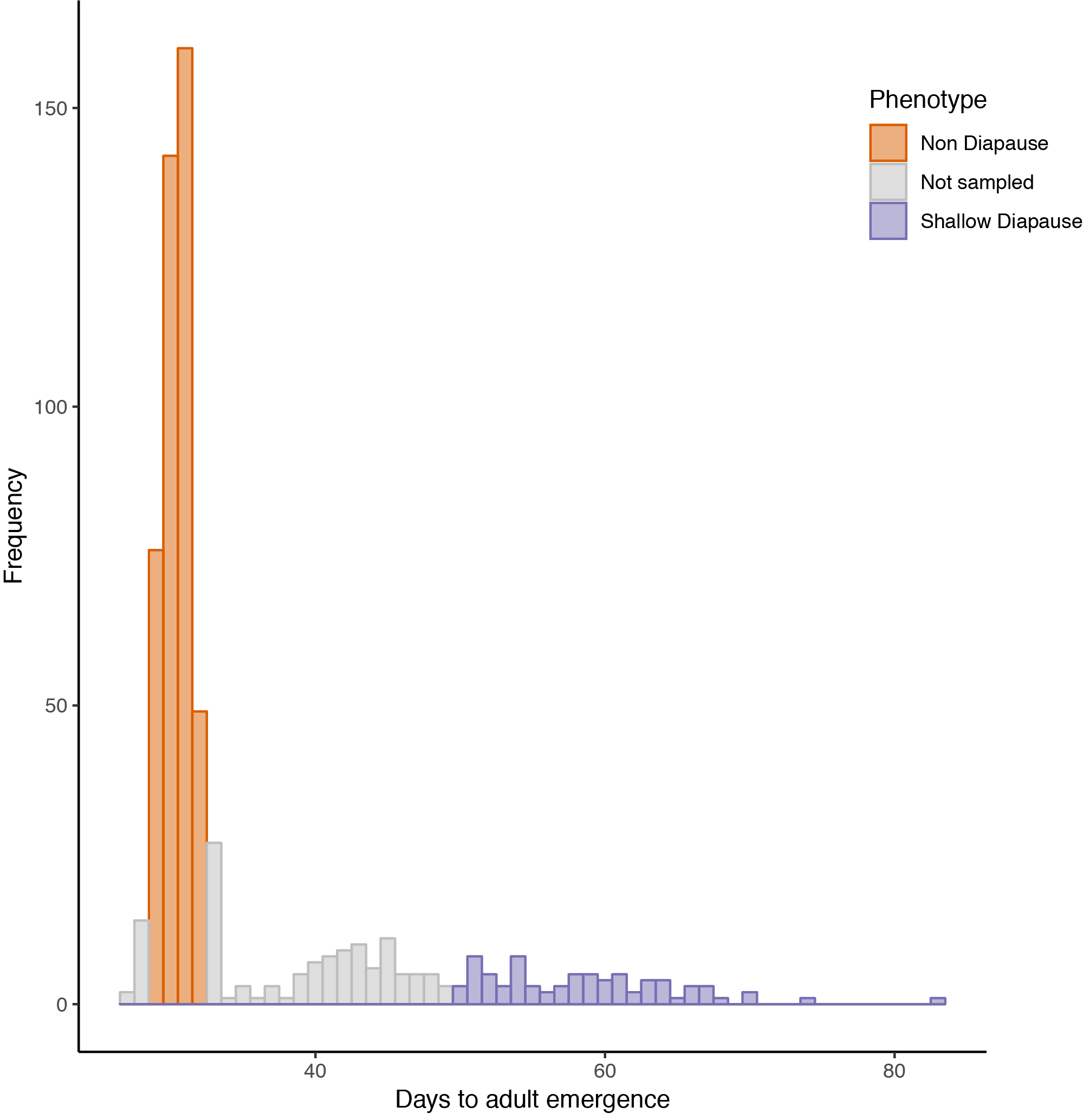

**Supporting Figure 2**: Histogram of hawthorn adult emergence times within the emergence experiment performed at the University of Notre Dame. Orange bins are individuals that were classed as ND, gray bins are individuals who were unsampled for this experiment due to uncertainty in their diapause class, and blue bins are individuals that were classed as SD.

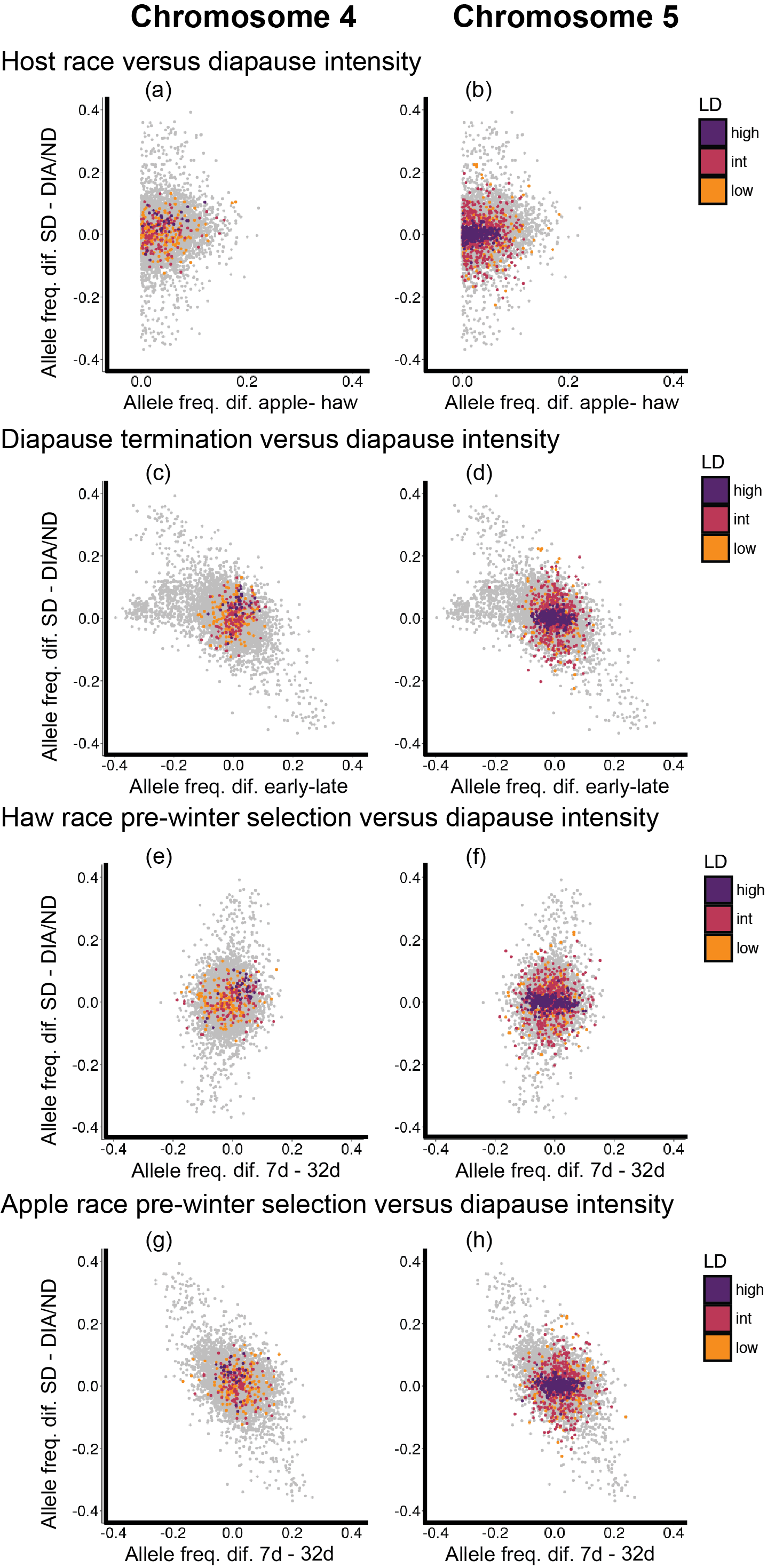

**Supporting figure 3:** Relationships between the strength of genetic associations (magnitude of allele frequency differences) with diapause phenotypes and host race differences, including diapause intensity (SD – (DIA+ND)) vs. (a,b) host race differences (apple host race – haw host race), (c,d) diapause termination (early – late), (e,f) hawthorn fly selection experiment (7 day – 32 day), and (g,h) apple fly selection experiment (7 day – 32 day). Light grey dots represent all 7,265 genotyped SNPs, while orange dots are low LD loci, red dots are intermediate LD loci, and purple dots are high LD loci on chromosome 4 (1^st^ column; a, c, e, g) and chromosome 5 (2^nd^ column; b, d, f, h). All loci were polarized to the allele at the highest frequency in the hawthorn fly population at Grant, MI; thus, host race differences in allele frequency were all positive.

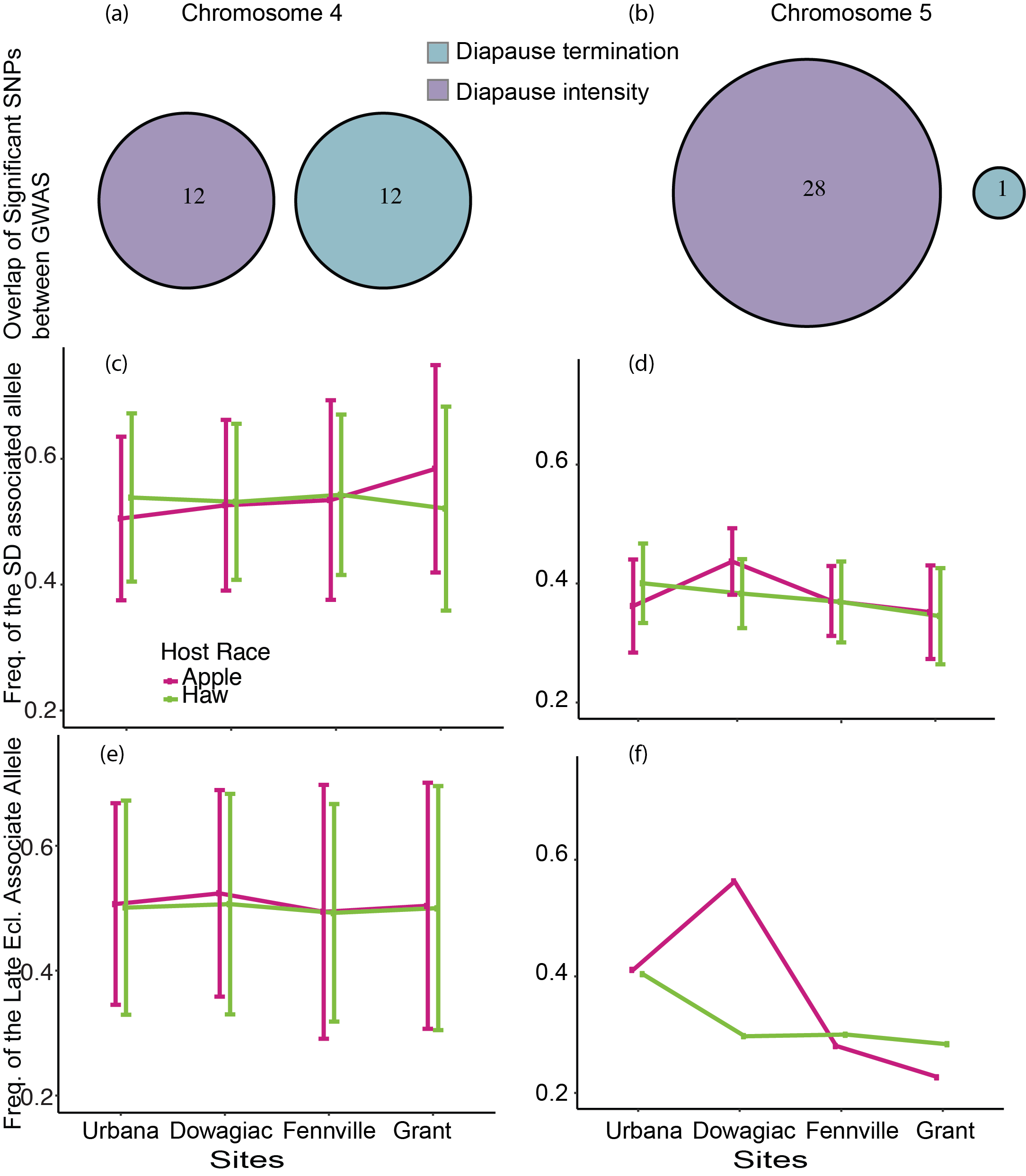

**Supporting figure 4:** (a,b) Overlap between and mean allele frequencies of loci significantly associated with diapause intensity and diapause termination on chromosomes 4 and 5. Venn Diagrams display overlap between loci significantly associated with diapause intensity and those associated with diapause termination. It should again be noted that the variation associated with diapause intensity on chromosome 5 may be do to unbalanced sex ratios in the samples genotyped for this experiment. The middle row (c,d) of line plots displays clinal variation in mean allele frequency with loci significantly associated with diapause intensity for chromosomes 4 and 5. Genetic loci were polarized such the allele associated with the SD phenotype was ‘counted.’ The bottom row of line plots (e,f) displays clinal variation in mean allele frequency with loci significantly associated with diapause intensity. There are no error bars in panel f because there was only one loci on chromosome significantly associated with diapause termination on chromosome 5.
