## Supplemental ISunformation for "The genomics of trait combinations and their influence on adaptive divergence"

**Supporting Information**

**Supporting information 1 - Explanation of the Bayesian Sparse Linear Mixed model**

The BSLMM consists of the following equation:

$$y= \mu+\boldsymbol{X}\beta+u+ \epsilon$$

where *y* is the vector of phenotypic measurements, μ is a vector of random effects modeling the infinitesimal effects of each locus, ***X*** is a *p* x *n* matrix of mean genotypes (*p* loci, *n* individuals), 𝜷 is the vector of ‘measurable’ effect sizes for each locus in **X**, *u* models random, infinitesimal effects, and 𝜖 is the error term.

**Supporting information 2 - Mean allele frequency estimates**

To measure clinal variation in SNPs significantly associated with either diapause termination or diapause intensity, mean allele frequencies were calculated at those loci within each host race at each sympatric site. The mean allele frequency was calculated by averaging the mean genotype scores for all individuals within a given host race at a given site for loci that were significantly associated with either diapause termination or diapause intensity.

**Supporting information 3 - Permutation tests**

Tests for allele frequency differences were performed using a nonparametric Monte Carlo approach between sample pools. This method established a null distribution of allele frequency differences at each SNP for a given pair of sample pool comparisons (e.g. early – late diapause termination, SD-ND/DIA, Apple-Haw, Grant-Urbana, etc.) by generating a set of 10,000 randomized allele frequency difference calculations for each locus. To do this, a pair of randomized samples were generated by combining the two sample pools (e.g. early – late, etc) into one and randomly drawing with replacement from that pool to create two new pools of the same size as the original, unpermuted pool. An allele frequency difference was then calculated between those two pools. Each locus with an empirical allele frequency difference that exceeded the upper 95% quantile of the null Monte Carlo distribution was taken as evidence the locus frequency was significantly different between the pools under study.

We also performed significance tests for linear regression analyses using a Monte Carlo resampling approach. When comparing responses from two experiments, a null distribution of 10,000 allele frequency difference correlations was calculated by first permuting a set of allele frequency differences within each comparison as above and then calculating a correlation coefficient between the resampled allele frequency differences. Empirical correlation coefficients that exceeded the abolsute upper 95% quantile of permuted correlation coefficients were taken as significant associations.

**Supporting information 4 - X-fold tests**

If sets of loci are strongly differentiated in two comparisons but at relatively uniform magnitudes (e.g., associated with diapause termination and initiation), the above correlation analysis may fail to detect a relationship. To account for this, we also implemented the X-fold test as described in (Chaturvedi et al. 2018). This approach tests whether more loci that expected by chance exhibit the same or opposite sign allele frequency differences for two comparisons (here, differences between eclosion bulks and differences between diapause intensity classes). To ask if the number of loci exhibiting sign concordance between the two GWA studies was greater than expected by chance we permuted whole genotypes and calculated new allele frequency differences for each GWA study to produce a null distribution of sign concordance. If the empirical number of loci with sign concordance for a given LD-chromosome group was greater the 95^th^ percentile of the null distribution, the level sign concordance was assumed to be greater than what would be expected by chance.
