## Supplemental Table 3 for "The genomics of trait combinations and their influence on adaptive divergence"

|  | chr 1 |  | chr 2 |  | chr 3 |  | | chr 4 | |  | | chr 5 | |  | | chr1-5 | |
| --- | --- | --- | --- | --- | --- | --- | --- | --- | --- | --- | --- | --- | --- | --- | --- | --- | --- |
| A) **Host Race** |  |  |  |  |  | |  | |  | |  | |  | |  | |  |
| All SNPs | 690 |  | 466 |  | 758 | |  | | 316 | |  | | 941 | |  | | 3171 |
| High LD | 208 |  | 81 |  | 176 | |  | | 35 | |  | | 296 | |  | | 796 |
| Int. LD | 383 |  | 324 |  | 438 | |  | | 117 | |  | | 467 | |  | | 1729 |
| Low LD | 99 |  | 61 |  | 144 | |  | | 164 | |  | | 178 | |  | | 646 |

**Supporting Table 3:** Total number of SNPs mapped to chromosomes and assigned to LD groups.
