## Supplemental Table 4 for "The genomics of trait combinations and their influence on adaptive divergence"

| Comparison | PGE mean estimate | PGE 95% Credible Interval |
| --- | --- | --- |
| SD vs. DIA | 0.19 | 0 – 0.79 |
| SD vs. ND | 0.42 | 0 – 0.94 |
| DIA vs. ND | 0.44 | 0 – 0.97 |

**Supporting Table 4:** The mean and 95% credible interval for the distribution of model estimates for the proportion of variances explained by the sparse effect term (PGE) generated from the BSLMM (Supporting information 1).
